# Interferon regulatory factor 3 upregulates the Treg recruitment factor CCL22 in response to double-stranded DNA in cancer cells

**DOI:** 10.1101/2022.03.08.483519

**Authors:** Jihyun G. Kim, Jocelyn V. Peña, Hannah P. McQueen, Lingwei Kong, Dina Michael, Pamela R. Cook

## Abstract

Cancer immunotherapy holds great promise for the treatment of solid tumors, but its effectiveness is hindered by the recruitment of regulatory T cells (Tregs), which inhibit anti-tumor immune responses. We report here that cytosolic dsDNA, a characteristic of many cancer cells, upregulates expression of the Treg-recruitment chemokine CCL22 in multiple types of malignant epithelial cells. We also identified that interferon regulatory factor 3 (IRF3) is a key regulator of CCL22 in response to dsDNA. Both IRF3 and NF-κB are activated downstream of the stimulator of interferon genes (STING), a primary effector protein responding to multiple cytosolic dsDNA sensors. IRF3 activation by STING triggers robust expression of type I interferons, which can boost anti-tumor immune responses. Thus, STING agonists have been used clinically to activate IRF3 during immunotherapy. However, STING activation in some cases is reported to paradoxically foster a pro-tumor, immunosuppressive environment. Our finding that IRF3 regulates CCL22 in response to dsDNA suggests a possible mechanism contributing to STING-mediated immunosuppression. In addition, we found that cultured cancer cells appear able to evolve mechanisms to co-opt nucleic acid sensing pathways to upregulate CCL22, suggesting that these pathways may contribute to acquired immune evasion in tumors with increased cytosolic dsDNA.

## Introduction

Recruitment of regulatory T cells (Tregs) to the tumor microenvironment remains one of the most significant barriers to successful immunotherapy for epithelial solid tumors. A primary mechanism of Treg recruitment is the chemokine CCL22 (MDC), which binds the CCR4 receptor expressed preferentially on T cell subsets including type 2 helper T cells and Tregs (1-4). The biological significance of increased CCL22 in the tumor microenvironment has been documented in numerous studies, and the majority report an association between increased CCL22 and poor prognoses (2,5-29).

CCL22 was first characterized in macrophages, but several reports have shown that CCL22 can also be expressed by various types of malignant cells (6,20,25,30-34). Additional studies have shown that CCL22 expressed directly from explanted malignant cells recruits Tregs and Fox3+ cells *in vivo* (15,30). Blocking the interaction between CCL22 and its receptor with the anti-CCR4 antibody Mogamulizumab was found to deplete Tregs during treatment of refractory adult T cell leukemia/lymphoma and cutaneous T cell lymphoma, prompting trials to test its effectiveness in advanced solid tumors, with mixed results (reviewed in 2). A potential drawback to a CCR4 blockade is that the chemokine CCL17 also binds CCR4 and is reported to have nonredundant and even opposing functions to CCL22, with CCL17 tending to promote inflammatory responses and CCL22 inducing immune tolerance (10,reviewed in 35). It has thus been suggested that antagonism of CCL22, rather than CCR4, may be a superior strategy for anti-cancer therapy (32). Pharmacological inhibition of CCL22 expression may also be useful and would be facilitated by a better understanding of its regulation in cancer cells.

Due to the initial discovery of CCL22 in macrophages and a later association with atopic dermatitis, much work has focused on elucidating the regulatory networks of CCL22 in myeloid lineages and keritonocytes. These studies have collectively suggested that the regulatory landscape of CCL22 is context-dependent and may differ with both cell type and species. For example, it was reported that CpG-oligodeoxynucleotides (CpG-ODN) strongly enhanced CCL22 expression in murine dendritic cells (10), but another study concluded that CpG-ODN inhibited CCL22 in tumor-associated murine dendritic cells (17). CpG-ODN were also found to repress CCL22 across a range of murine bulk tumor samples (17) and in a murine asthma model (36) but increased CCL22 in cell isolates from human ovarian tumors (17). Another example of cell-specific regulation of CCL22 is its relationship with interferon-gamma, with studies reporting that it increased CCL22 in human keratinocytes (37-39), had no effect on CCL22 in human fibroblasts (40) or human airway smooth muscle cells (41), inhibited CCL22 in monocytes and macrophages (42), and was inversely correlated with CCL22 in T cells (43). These and other studies demonstrate that our understanding of CCL22 regulation remains incomplete, particularly with respect to human cancers.

We have found that CCL22 is upregulated in several human malignant epithelial cell lines in response to cytosolic double-stranded DNA (dsDNA). Healthy normal cells restrict DNA to the nucleus, and its presence in the cytosol is indicative of pathogens or aberrantly localized self-DNA, which can occur in cancer and certain autoimmune diseases. Accumulation of self DNA in the cytosol of cancer cells can be due to genomic instability, damaged mitochondria, and reactivated transposable elements. A variety of nucleic acid sensors detect this DNA and initiate a signaling cascade through the effector protein stimulator of interferon genes (STING), triggering proinflammatory innate and adaptive immune responses via STING-mediated activation of the transcription factors NF-κB and interferon regulatory factor 3 (IRF3). One of the most well-described sensors is the cyclic GMP-AMP synthase (cGAS) (reviewed in 44,45). In addition to detecting cytosolic dsDNA, cGAS reportedly binds cytosolic RNA:DNA hybrids, and recent reports suggest it may also respond to LINE-1 cDNA (46-49,reviewed in 50). Upon nucleic acid binding, cGAS synthesizes the second messenger cyclic dinucleotide 2’3’-cGAMP, which in turn binds STING, prompting oligomerization and a conformational change that allows phosphorylation of STING on S366 by TANK-binding kinase 1 (TBK1). The interaction between STING and TBK1 brings about the phosphorylation of IRF3 by TBK1 (51-53), initiating IRF3-mediated upregulation of type I interferons. Type I interferons mediate robust inflammatory responses, thus the use of STING agonists in immunotherapy has gained widespread interest for their potential to intensify therapeutic anti-tumor responses. However, STING activation in some cases has been reported to paradoxically contribute to pro-tumor, immunosuppressive environments (50,54-58). The ability of dsDNA and STING agonists to robustly upregulate CCL22 may be a potential mechanism of STING-mediated immune suppression. In the present study, we used a combination of reverse genetic and biochemical approaches to investigate the specific contributions of each downstream STING pathway, IRF3 and NF-κB, to dsDNA-mediated upregulation of CCL22.

## Results

### The chemokine CCL22 is strongly induced in tumor cells by cytoplasmic dsDNA

During our investigation of LINE-1 re-activation in epithelial cancer cells, we found that dsDNA transfected into cells robustly increased CCL22 expression, while a mock control (transfection reagent alone) had no effect (Fig. 1A). The ability of cytosolic dsDNA to elicit robust immune signaling is well-known, but the magnitude of CCL22 upregulation was so great that we first considered the possibility that our DNA might be contaminated with endotoxin, notwithstanding the use of endotoxin-free DNA purification procedures. Treatment with DNA alone in the absence of transfection reagent did not increase CCL22, confirming that DNA entry into cells was required for CCL22 upregulation (Fig. S1).

**Figure 1.**
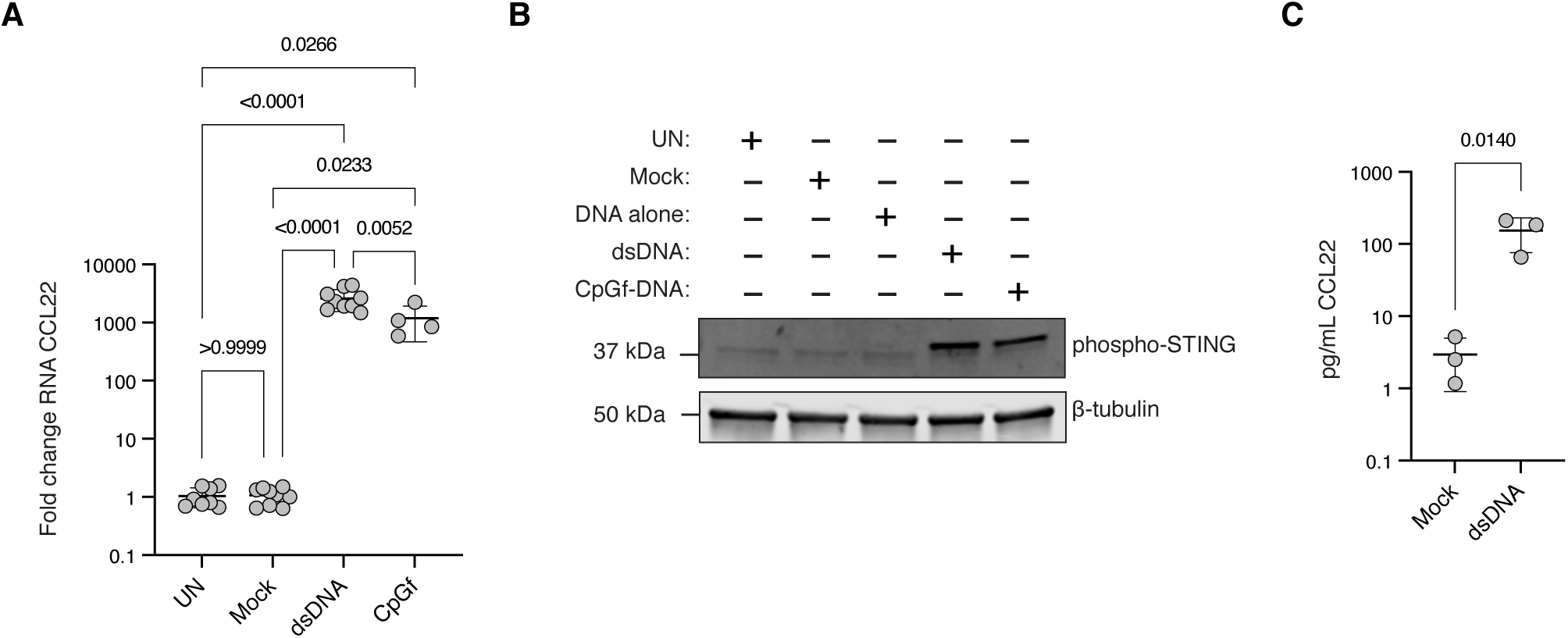
Double-stranded DNA increases CCL22 in HeLa cells. *A*, HeLa cells were untreated (UN) or treated with a mock control, dsDNA (2 μg/mL), or CpG-free (CpGf) dsDNA (2 μg/mL) using *Trans*IT-LT1 at a 1:2 ratio and harvested 48 hours after transfection. Resulting levels of CCL22 mRNA are shown. Each data point represents an independent experiment with values derived from three technical replicates. Significance testing was performed with a one-way ANOVA and Tukey’s pairwise comparison; error bars represent standard deviations. *B*, HeLa cells were transfected as described in *A* and harvested after 48 hours. Lysates (20 μg) were separated with SDS-PAGE and probed for phospho-STING (S366) and beta-tubulin. The image shown is representative of at least three independent experiments. *C*, HeLa cells were transfected as in *A*. Media was harvested 48 hours after treatment and analyzed with an ELISA for CCL22 protein. Each data point represents an independent experiment with values derived from three technical replicates. Significance testing was performed with an unpaired, one-tailed t test; error bars represent standard deviations.

We next sought to determine whether CpG motifs in our DNA might be responsible for CCL22 upregulation. Previous studies investigating the effects of CpG oligonucleotides on CCL22 expression, taken together, suggest that CpGs can both inhibit and stimulate expression of CCL22, perhaps in a cell-type and species-specific manner (10,17). In theory, transfected CpG-containing dsDNA could be digested intracellularly to produce CpG-containing single-stranded DNA (59,reviewed in 60), a potent ligand of endosomal toll like receptors 9 (TLR9). However, we found that CpG-free dsDNA was only slightly less efficient at upregulating CCL22, indicating that CpG-mediated activation of TLR9 was not the primary factor driving CCL22 expression in our system and that another dsDNA-sensing pathway was involved (Fig. 1A).

STING is the primary effector protein for multiple cytosolic dsDNA sensors, including cGAS. In order to confirm that dsDNA activated STING in our cells, we probed for phosphorylated STING S366, which is targeted by TBK1 in response to dsDNA (51-53). Figure 1B shows that dsDNA, with or without CpGs, increased phosphorylation of STING S366 compared to untreated cells or cells treated with a mock transfection reaction (no DNA) or DNA alone (no transfection reagent). CCL22 protein was also significantly increased in the media of cells treated with dsDNA, confirming that increased CCL22 mRNA resulted in increased protein, and that the protein was secreted (Fig. 1C).

To determine whether dsDNA would increase CCL22 expression in other epithelial cancer types, five additional cell lines were tested. Two of the five were not amenable to transfection, and the other three, MCF7, JEG-3, and HCT 116, all increased CCL22 expression in response to dsDNA to statistically significant levels (Fig. 2A-C). These findings suggest that upregulation of CCL22 in response to dsDNA may be a common finding across multiple types of epithelial cancer cells. In order to confirm that DNA was effectively delivered to each cell line, parallel experiments were performed using a GFP expression plasmid, and GFP was subsequently visualized in live cells (Fig. 2D). Although it is possible that slightly reduced DNA delivery to MCF7 cells may have contributed to the reduced level of CCL22 upregulation in those cells, the relatively small decreases in DNA delivery to JEG-3 and HCT 116 cells is unlikely to account for the large differences in their response compared to HeLa and MCF7 cells.

**Figure 2.**
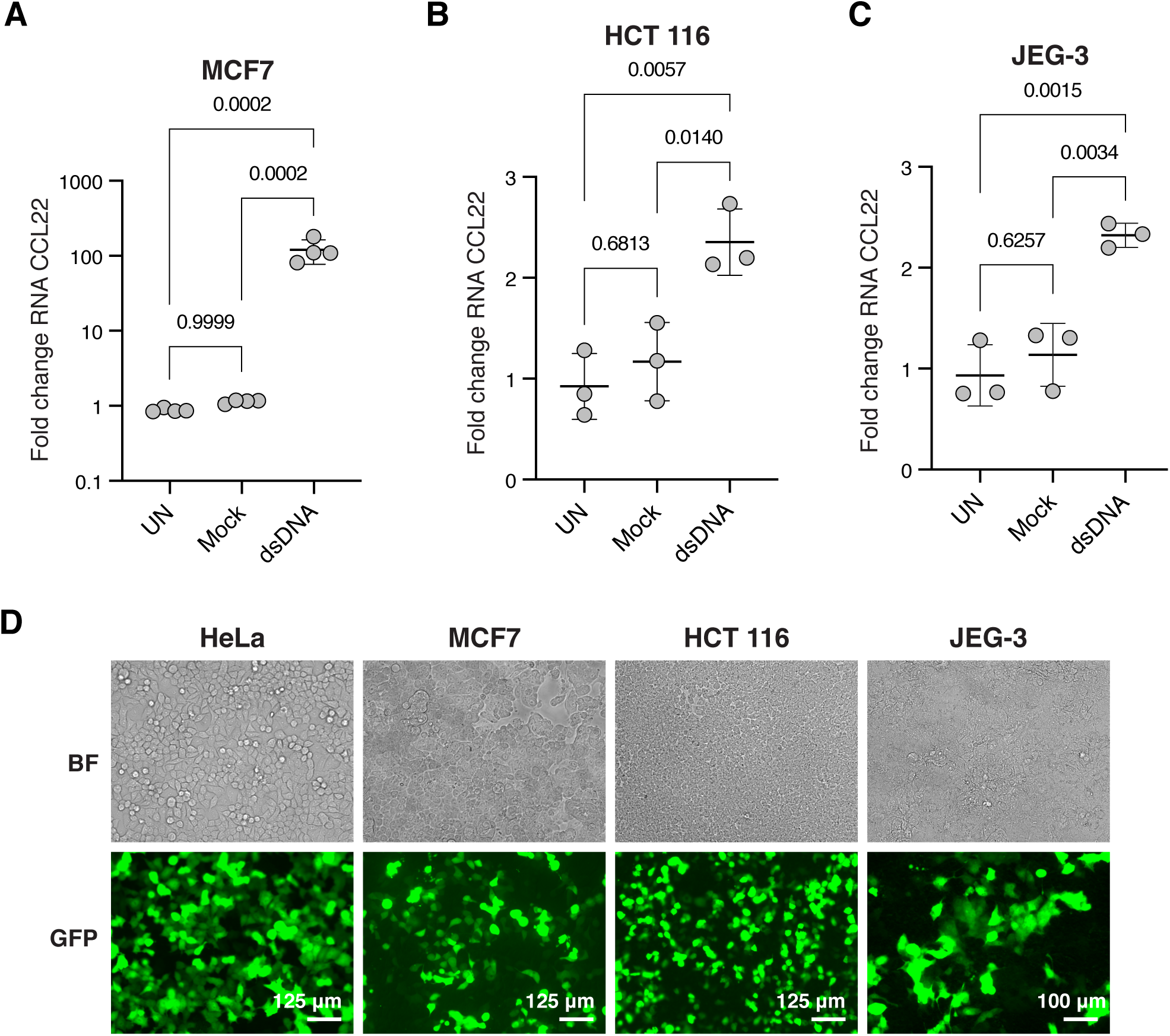
Double-stranded DNA increases CCL22 in multiple types of epithelial cancer cells. *A-C*, MCF7 (*A*), HCT 116 (*B*), and JEG-3 (*C*) cells were untreated (UN) or treated with a mock control, dsDNA (2 μg/mL) using TransfeX at a 1:4 ratio (MCF7) or *Trans*IT-LT1 (HCT 116, 1:4 ratio; JEG-3, 1:3 ratio). Cells were harvested 48 hours after transfection. Resulting levels of CCL22 mRNA are shown. Each data point represents an independent experiment with values derived from three technical replicates. Significance testing was performed with a one-way ANOVA and Tukey’s pairwise comparison; error bars represent standard deviations. *D*, Cells were transfected as in *A-C* in parallel experiments using a GFP expression plasmid (2 μg/mL with transfection reagents and ratios indicated in *A-C*) and imaged 48 hours after transfection. Brightfield (BF) shows the confluency of cells in the same field of view as GFP.

### Pharmacological activation of STING upregulates CCL22

To determine whether direct STING activation also upregulated CCL22, cells were treated with a stabilized analog of the canonical STING agonist 2’3’-cGAMP, 2’3’-cGAM(PS)_2_(Rp/Sp). Consistent with our findings using dsDNA, direct activation of STING with 2’3’-cGAM(PS)_2_(Rp/Sp) increased CCL22 expression in HeLa cells (Fig. 3A), but at lower levels than observed after treatment with dsDNA. Preliminary experiments had indicated optimal time points after treatment to capture maximum levels of CCL22 upregulation in response to 2’3’-cGAM(PS)_2_(Rp/Sp) (24 hours post treatment) and dsDNA (48 hours post treatment). Thus, the difference in CCL22 expression in response to each stimulus was not due to different treatment times, as cells were harvested at the time point of maximum CCL22 expression for each treatment type. Figure 3B confirms that 2’3’-cGAM(PS)_2_(Rp/Sp) induces STING phosphorylation, with peak phosphorylation occurring at approximately 6 hours post treatment and declining almost to baseline by 24 hours (Fig. 3B), compared to STING phosphorylation following treatment with dsDNA, which was still apparent after 48 hours (Fig. 1B). In MCF7 cells, 2’3’-cGAM(PS)_2_(Rp/Sp) upregulated CCL22 (Fig. 3C) to approximately the same levels as dsDNA (Fig. 2A). We did not test the effect of 2’3’-cGAM(PS)_2_(Rp/Sp) in the other cell lines, as CCL22 upregulation in those cells was already much lower than in HeLa and MCF7 cells.

**Figure 3.**
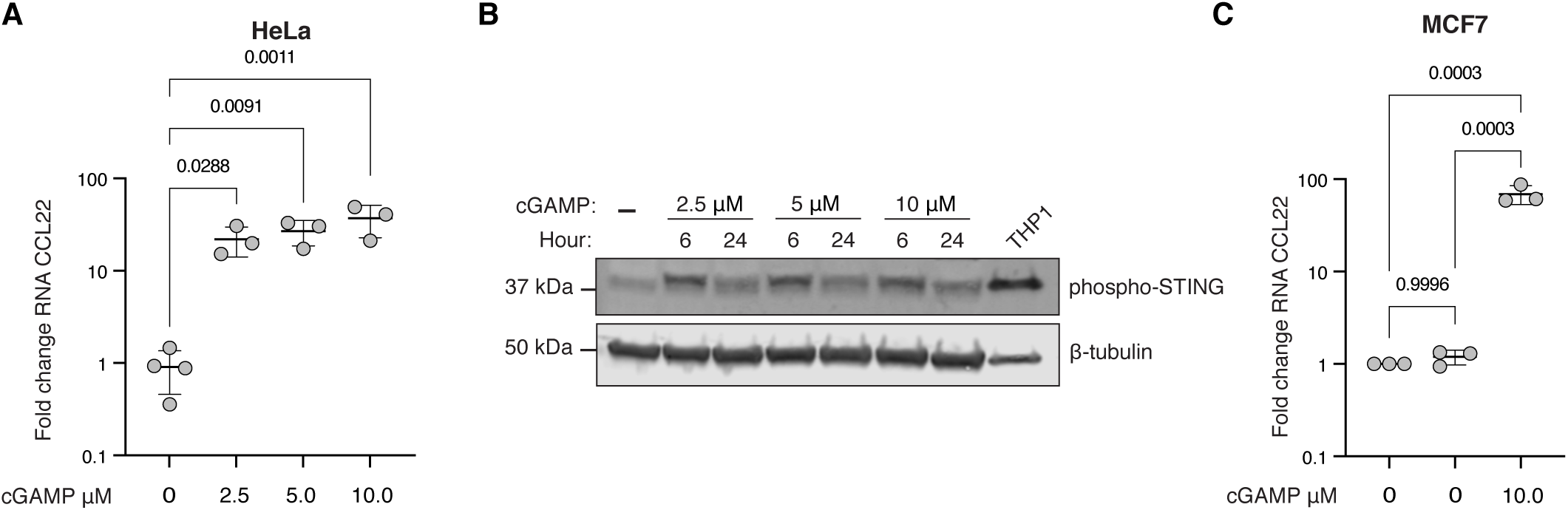
STING agonist 2’3’-cGAM(PS)_2_(Rp/Sp) upregulates CCL22. *A*, HeLa cells were transfected with a mock control or 10 μM 2’3’-cGAM(PS)_2_(Rp/Sp) with *Trans*IT-LT1 at a 1:1 μg/μL ratio and harvested 24 hours post transfection for RTqPCR. Resulting levels of CCL22 are shown relative to untreated cells. Each data point represents an independent experiment with values derived from three technical replicates. Significance testing was performed with a one-way ANOVA and Dunnett’s multiple comparison test; error bars represent standard deviations. *B*, HeLa cells were transfected with a mock control or indicated concentrations of 2’3’-cGAM(PS)_2_(Rp/Sp) with *Trans*IT-LT1 at a 1:1 μg/μL ratio and harvested at indicated timepoints. Lysates (60 μg) and THP-1 positive control (15 μg) were resolved with SDS-PAGE and probed for pSTING and beta-tubulin. *C*, MCF7 cells were transfected with a mock control or 10 μM 2’3’-cGAM(PS)_2_(Rp/Sp) with TransfeX at a 1:1 μg/μL ratio and harvested 24 hours post transfection for RTqPCR. Resulting levels of CCL22 are shown relative to untreated cells. Each data point represents an independent experiment with values derived from three technical replicates. Significance testing was performed with a one-way ANOVA and Tukey’s pairwise comparison; error bars represent standard deviations.

### Inhibition of TBK1/IKKε abrogates dsDNA-mediated activation of CCL22

STING phosphorylation, as well as phosphorylation and activation of NF-κB and IRF3, depend on TBK1 and/or its homolog, I-kappa-B kinase (IKK) epsilon (IKKε) (61-64). Pretreatment of cells with MRT67307, a reversible inhibitor of TBK1/IKKε, led to a robust and dose-dependent decrease in dsDNA-mediated CCL22 upregulation in both HeLa and MCF7 cells, without affecting transfection efficiency in either cell line (Fig. 4A-D). MRT67307 also inhibited CCL22 upregulation in response to 2’3’-cGAM(PS)_2_(Rp/Sp) in both HeLa and MCF7 cells (Fig. 4E-F). Of note, although MRT67307 is reported by the supplier (Invivogen) to specifically inhibit IRF3 with no effect on NF-κB, this statement is based in part on testing by the supplier that relied on activating NF-κB with RNA hairpin ligands for the retinoic acid-inducible gene I (RIG-1), which activates NF-κB via the IKKα/IKKβ pathway, thus avoiding dependence on TKB1/IKKε. However, in the context of STING-mediated activation of NF-κB, multiple studies have indicated the involvement of TBK1 and/or IKKε (61-64), and TKB1/IKKε have also been implicated in phosphorylation and activation of the canonical NF-κB subunit RELA/p65 (65,reviewed in 66). Our testing of MRT67307 showed that although it had no observable effect on RELA/p65 phosphorylation in response to the two well-known NF-κB activators tumor necrosis factor alpha (TNFα) and phorbol 12-myristate 13-acetate (PMA), (Fig. S2A), MRT67307 did reduce phosphorylation of RELA/p65 in response to 2’3’-cGAM(PS)_2_(Rp/Sp) and slightly reduced phosphorylation in response to dsDNA (Fig. S2B). These results are consistent with a previous study showing that MRT67307 slightly reduced p65 phosphorylation in response to the STING agonist DMXAA (63). We also confirmed that MRT67307, as expected, inhibited phosphorylation of IRF3 (S386) in response to both 2’3’-cGAM(PS)_2_(Rp/Sp) and dsDNA (Fig. S2C). To delineate the relative contributions of each downstream STING pathway, NF-κB and IRF3, to dsDNA-mediated activation of CCL22, we used RNAi against each pathway.

**Figure 4.**
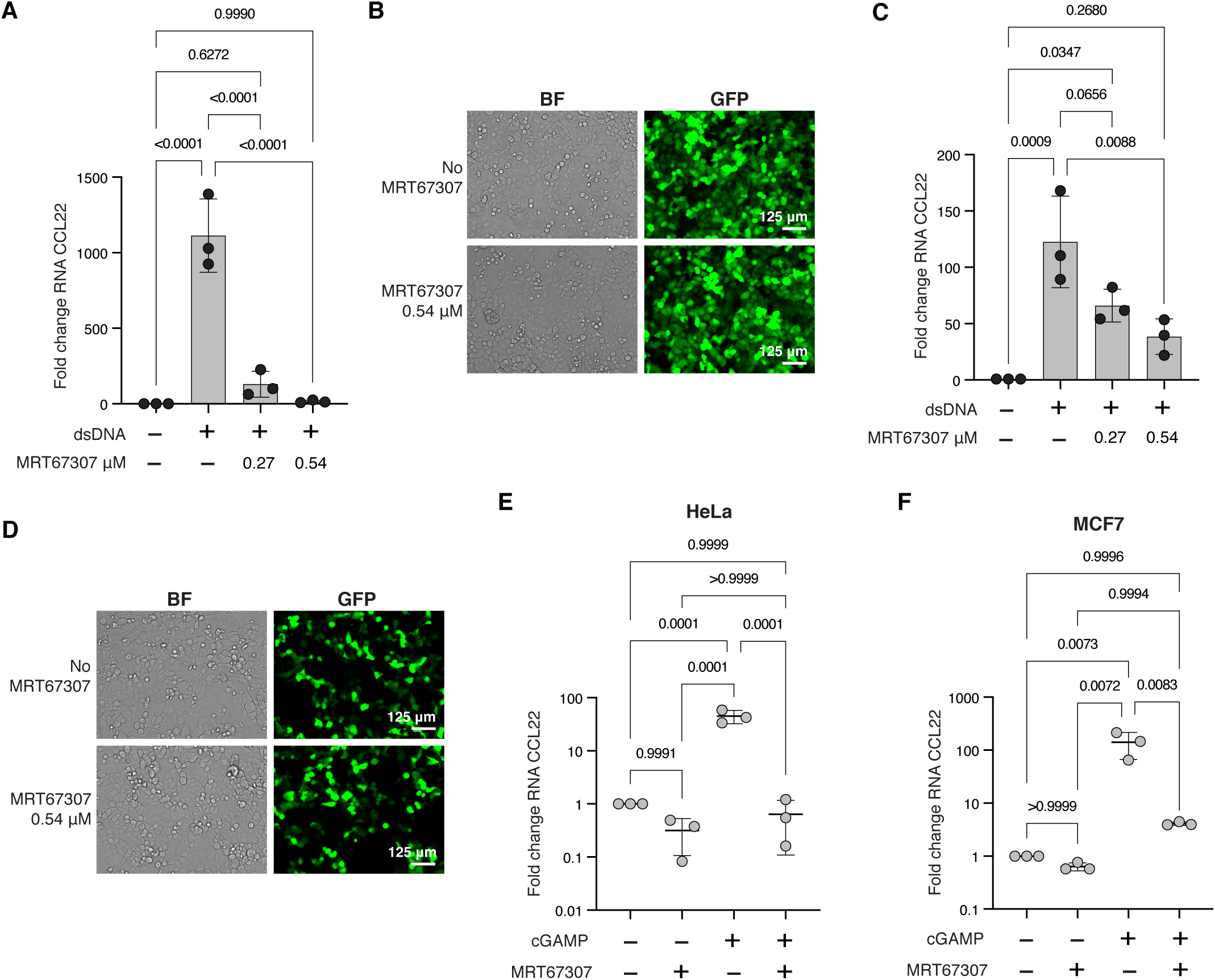
Inhibition of TBK1 and IKKε inhibit upregulation of CCL22 by dsDNA and STING agonist activation. *A*, HeLa cells were seeded in 12-well plates to achieve ∼ 65% confluency in 24 hours, then cells were treated with either the TBK1/IKKε inhibitor MRT67307 at concentrations indicated in the figure or a water vehicle control for approximately 1.5 hours prior to transfection with either a mock control or dsDNA (2 μg/mL) with *Trans*IT-LT1 at a 1:2 ratio. Cells were harvested 48 hours after transfection for RTqPCR. Resulting levels of CCL22 mRNA are shown. Each data point represents an independent experiment with values derived from three technical replicates. Significance testing was performed with a one-way ANOVA and Tukey’s pairwise comparison; error bars represent standard deviations. *B*, HeLa cells were treated with MRT67307 as described in *A*, then transfected in parallel experiments using a GFP expression plasmid (2 μg/mL, *Trans*IT-LT1 1:2 ratio) and imaged 48 hours after transfection. Brightfield (BF) shows the confluency of cells in the same field of view as GFP. *C*, MCF7 cells were seeded in 12-well plates to achieve ∼ 50% confluency in 24 hours, then cells were treated with either the TBK1/IKKε inhibitor MRT67307 at concentrations indicated in the figure or a water vehicle control for approximately 1.5 hours prior to transfection with either a mock control or dsDNA (2 μg/mL) with TransfeX at a 1:4 ratio. Cells were harvested 48 hours after transfection for RTqPCR. Resulting levels of CCL22 mRNA are shown. Each data point represents an independent experiment with values derived from three technical replicates. Significance testing was performed with a one-way ANOVA and Tukey’s pairwise comparison; error bars represent standard deviations. *D*, MCF7 cells were treated with MRT67307 as described in *C*, then transfected in parallel experiments using a GFP expression plasmid (2 μg/mL, TransfeX, 1:4 ratio) and imaged 48 hours after transfection. Brightfield (BF) shows the confluency of cells in the same field of view as GFP. *E-F*, HeLa cells (*E*) and MCF7 cells (*F*) were treated with 0.54 μM of the TBK1/IKKε inhibitor MRT67307 or a water vehicle control for approximately 1.5 hours prior to transfection with 10 μM 2’3’-cGAM(PS)_2_(Rp/Sp) using *Trans*IT-LT1 (HeLa) or TransfeX (MCF7) at a 1:1 μg/μL ratio or the respective mock controls. Cells were harvested 24 hours after transfection, and RTqPCR was performed. Resulting fold change of CCL22 is shown. Each data point represents an independent experiment with values derived from three technical replicates. Significance testing was performed with a one-way ANOVA and Tukey’s pairwise comparison; error bars represent standard deviations.

### NF-κB contributes to CCL22 upregulation by dsDNA

Previous studies investigating the regulation of CCL22 have identified a role for NF-κB (67-72). However, we found that treatment of HeLa cells with the NF-κB activators TNFα (Fig. 5A) and PMA (Fig. 5B) induced CCL22 expression at substantially lower levels than treatment with dsDNA. We therefore used RNAi against the canonical NF-κB subunit RELA/p65 to determine its contribution to CCL22 upregulation within the context of cytosolic dsDNA activation of STING. Five shRNA constructs from the Sigma MISSION collection were initially tested, and two (p65-1 and p65-2) that effectively knocked down RELA/p65 at the transcript (Fig. 5C) and protein levels (Fig. 5D) were chosen. Both p65-1 and p65-2 reduced CCL22 upregulation in response to dsDNA, but only p65-1 resulted in a statistically significant result (Fig. 5E), consistent with the greater degree of knockdown from p65-1. Parallel transfections with a GFP expression plasmid showed that decreased CCL22 expression in the shRNA cell lines was not due to decreased DNA delivery to those cells (Fig. 5F).

**Figure 5.**
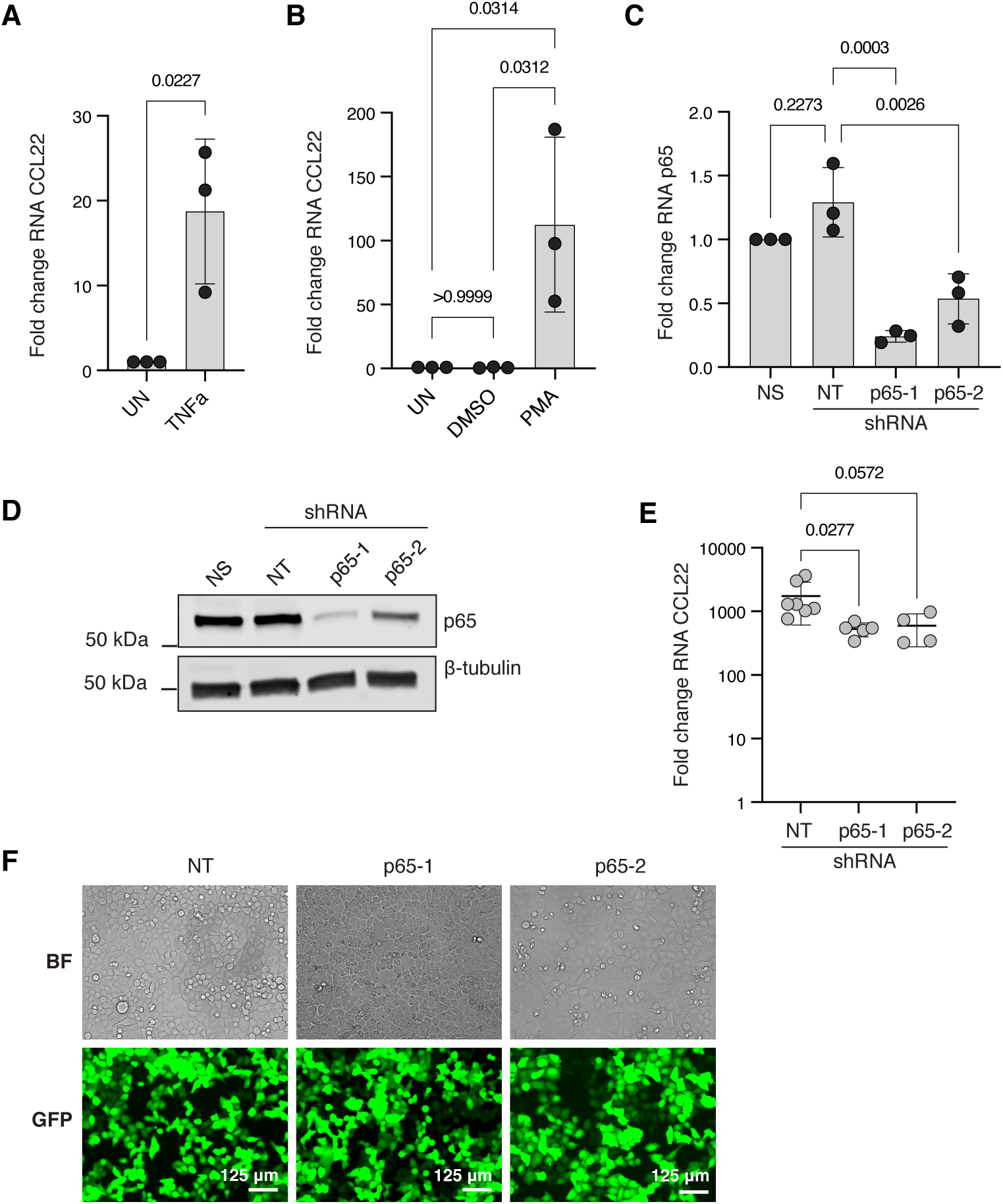
NF-κB contributes to CCL22 upregulation. *A*, HeLa cells were seeded in 12-well plates to achieve ∼ 65% confluency in 24 hours, at which time cells were treated with either 8 ng/mL TNFα or a water vehicle control. Cells were harvested 24 hours after treatment, and RTqPCR was performed. Resulting levels of CCL22 mRNA are shown. Each data point represents an independent experiment with values derived from three technical replicates. Significance testing was performed with an unpaired, two-tail t test; error bars represent standard deviations. *B*, HeLa cells were seeded and grown as for *A* and treated with either 10 ng/mL PMA or a 0.0004% DMSO vehicle control. Significance testing was performed with a one-way ANOVA and Tukey’s pairwise comparison; error bars represent standard deviations. *C*, HeLa cells expressing no shRNA (NS) or stably expressing a non-targeting shRNA (NT) or shRNAs against RELA/p65 (p65-1 or p65-2) were harvested for RTqPCR; levels of RELA/p65 RNA are shown relative to untreated (NS) cells. Each data point represents an independent experiment with values derived from three technical replicates. Significance testing was performed with a one-way ANOVA and Tukey’s pairwise comparison; error bars represent standard deviations. *D*, Lysates (20 μg) from HeLa cells carrying no shRNA (NS), non-targeting shRNA (NT) or shRNAs against RELA/p65 (p65-1 or p65-2) were resolved using SDS-PAGE and probed for RELA/p65 and beta-tubulin. The image shown is representative of at least three independent experiments. *E*, HeLa cells described in *C* and *D* were transfected with a mock control or dsDNA (2 μg/mL) with *Trans*IT-LT1 at a 1:2 ratio and harvested after 48 hours for RTqPCR. Resulting fold change of CCL22 mRNA is shown relative to the mock control for each individual cell line. Each data point represents an independent experiment with values derived from three technical replicates. Significance testing was performed with a one-way ANOVA and Dunnett’s pairwise comparison of each shRNA group to the control non-targeting group; note that this analysis was performed alongside the IRF3 shRNA groups from Figure 6C in order that all non-targeting control experiments be included; error bars represent standard deviations. *F*, HeLa cells carrying the shRNAs described above were transfected as in *E* in parallel experiments using a GFP expression plasmid (2 μg/mL, *Trans*IT-LT1 1:2 ratio) and imaged 48 hours after transfection. Brightfield (BF) shows the confluency of cells in the same field of view as GFP.

### IRF3 is required for CCL22 upregulation by dsDNA

The relatively weak effect of NF-κB knockdown compared to treatment with MRT67307, which also inhibits IRF3, suggested a predominant role for IRF3 in CCL22 upregulation. As before, two targeting shRNAs from a pool of five from the MISSION library were chosen that effectively knocked down IRF3 at both the transcript (Fig. 6A) and protein levels (Fig. 6B). Consistent with results using MRT67307, shRNA knockdown of IRF3 almost completely eliminated CCL22 upregulation in response to dsDNA, reducing the mean fold changes to 15.7 and 32.6 in the IRF3-1 and IRF3-2 shRNA samples, respectively (Fig. 6C). Parallel transfections with a GFP expression plasmid confirmed that differences were not due to variation in DNA delivery between the knock down lines (Fig. 6D).

**Figure 6.**
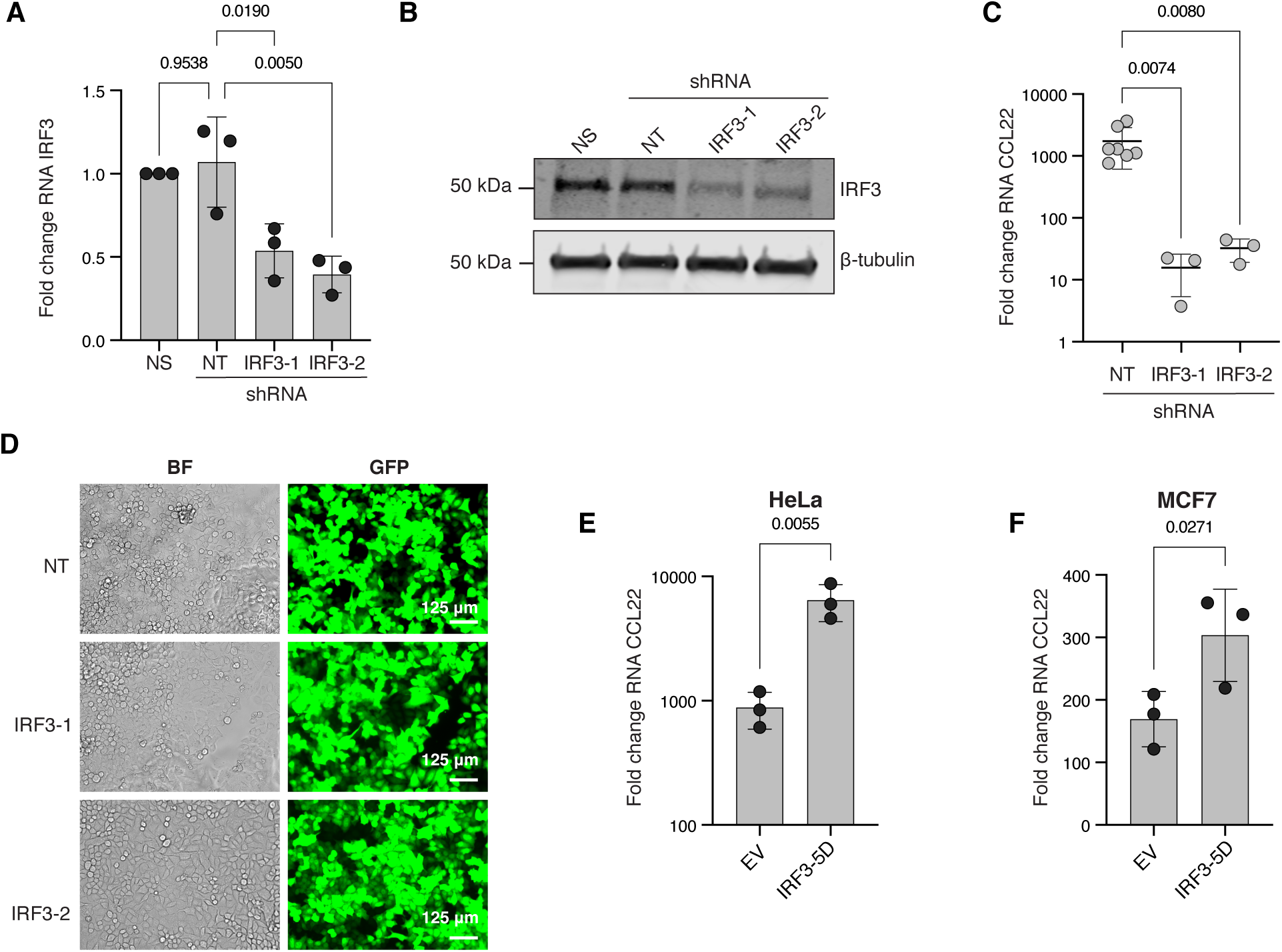
IRF3 is indispensable for CCL22 upregulation in response to dsDNA. *A*, HeLa cells expressing no shRNA (NS) or stably expressing a non-targeting shRNA (NT) or shRNAs against IRF3 (IRF3-1 or IRF3-2) were harvested for RTqPCR; levels of IRF3 RNA are shown relative to untreated (NS) cells. Each data point represents an independent experiment with values derived from three technical replicates. Significance testing was performed with a one-way ANOVA and Tukey’s pairwise comparison; error bars represent standard deviations. *B*, Lysates (20 μg) from HeLa cells carrying no shRNA (NS), non-targeting shRNA (NT) or shRNAs against RELA/p65 (p65-1 or p65-2) were resolved using SDS-PAGE and probed for RELA/p65 and beta-tubulin. The image shown is representative of at least three independent experiments. *C*, HeLa cells described in *A* and *B* were transfected with a mock control or dsDNA (2 μg/mL) with *Trans*IT-LT1 at a 1:2 ratio and harvested after 48 hours for RTqPCR. Resulting fold change of CCL22 mRNA is shown relative to the mock control for each individual cell line. Each data point represents an independent experiment with values derived from three technical replicates. Significance testing was performed with a one-way ANOVA and Dunnett’s pairwise comparison of each shRNA group to the control non-targeting group; note that this analysis was performed alongside the RELA/p65 shRNA groups from Figure 5E in order that all non-targeting control experiments be included; error bars represent standard deviations. *D*, HeLa cells carrying the shRNAs described above were transfected as in *C* in parallel experiments using a GFP expression plasmid (2 μg/mL, *Trans*IT-LT1, 1:2 ratio) and imaged 48 hours after transfection. Brightfield (BF) shows the confluency of cells in the same field of view as GFP. *E-F*, HeLa cells (*E*) and MCF7 cells (*F*) were transfected with a mock control or 2 μg/mL of empty plasmid (EV) or the constitutively active IRF3-5D with *Trans*IT-LT1 (HeLa) at a 1:2 ratio or TransfeX (MCF7) at a 1:4 ratio. Cells were harvested 48 hours after transfection, and RTqPCR was performed. Resulting fold change of CCL22 mRNA relative to the mock control is shown. Each data point represents an independent experiment with values derived from three technical replicates. Significance testing was performed with an unpaired, one-tailed t test; error bars represent standard deviations.

Given the effect of IRF3 RNAi on dsDNA-mediated upregulation of CCL22, a constitutively active IRF3 would be expected to further increase CCL22 above the level induced by the empty plasmid. The constitutively active IRF3-5D phosphomimetic has been well-characterized in the literature and carries five aspartic acid substitutions: S396D, S398D, S402D, T404D, and S405D (73-76). IRF3-5D significantly increased CCL22 in both HeLa (Fig. 6E) and MCF7 cells (Fig. 6F) above levels observed from the empty plasmid alone.

### Different strains of the same cell line differentially upregulate CCL22

Due to the magnitude of the effect of dsDNA on CCL22 expression in HeLa cells, we purchased new HeLa cells from the American Type Culture Collection (ATCC) to determine whether the same effect would be observed in those cells. Remarkably, CCL-2 HeLa cells from ATCC exhibited no significant increase of CCL22 expression in response to dsDNA (Fig. 7A). To confirm the authenticity of our original HeLa cells, both lines were sent to ATCC for short tandem repeat (STR) analysis, which showed a 100% match (Fig. 7B). We also confirmed that CCL-2 HeLa cells from ATCC were efficient at taking up dsDNA, as evidenced by GFP expression (Fig. 7C). To determine whether CCL-2 HeLa cells from ATCC activated STING in response to dsDNA, STING S366 phosphorylation was compared in both cell lines. Although STING phosphorylation was reduced in CCL-2 HeLa cells from ATCC, it was nonetheless detectable (Fig. 7D).

**Figure 7.**
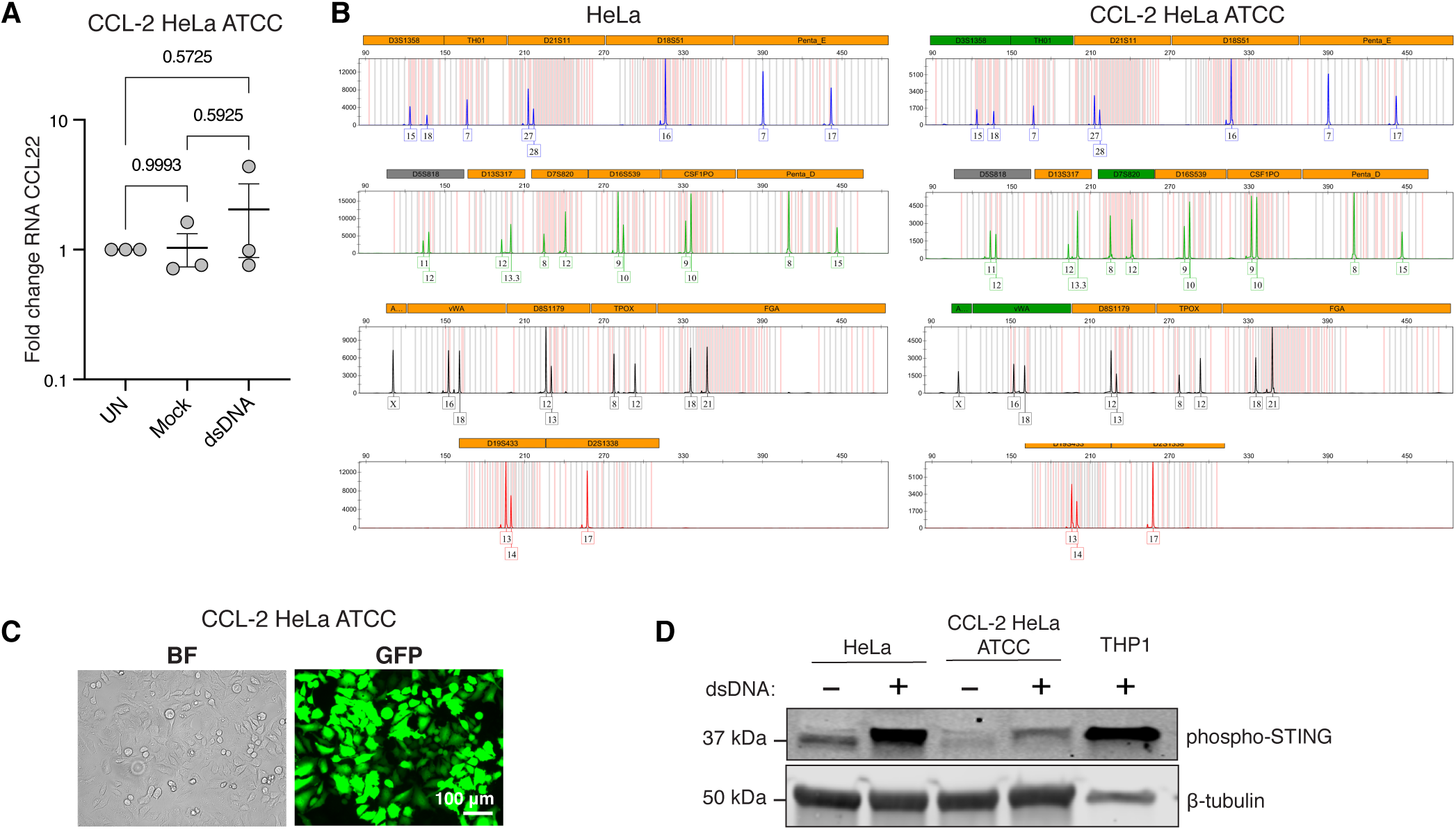
Two strains of HeLa cells differ dramatically in upregulation of CCL22 by dsDNA. *A*, CCL-2 HeLa cells from ATCC were untreated or transfected with a mock control or dsDNA (2 μg/mL) with *Trans*IT-LT1 at a 1:2 ratio and harvested after 48 hours for RTqPCR. Resulting fold change of CCL22 mRNA is shown. Each data point represents an independent experiment with values derived from three technical replicates. Significance testing was performed with a one-way ANOVA and Tukey’s pairwise comparison; error bars represent standard deviations. *B*, Samples of our original HeLa cell line and CCL-2 HeLa cells from ATCC were each sent to ATCC for authentication with STR profile analysis. Results are shown side-by-side. *C*, CCL-2 HeLa cells from ATCC were transfected as in *A* in parallel experiments using a GFP expression plasmid (2 μg/mL, *Trans*IT-LT1 ,1:2 ratio) and imaged 48 hours after transfection. Brightfield (BF) shows the confluency of cells in the same field of view as GFP. *D*, Indicated cell lines were transfected as described in *A* and harvested after 48 hours. Lysates (60 μg) and THP-1 positive control (15 μg) were separated with SDS-PAGE and probed for phospho-STING (S366) and beta-tubulin. The image shown is representative of at least three independent experiments.

### IRF3 activation in response to dsDNA does not correlate with STING phosphorylation in MCF7 cells

The lack of CCL22 upregulation in CCL-2 HeLa cells from ATCC, combined with reduced STING phosphorylation on S366, raised the question of whether the other cell lines exhibiting low levels of CCL22 upregulation, JEG-3 and HCT 116, also showed reduced STING phosphorylation. Figure 8A shows that neither JEG-3 nor HCT 116 cells had detectable levels of STING phosphorylation following treatment with dsDNA. More surprising however was the lack of STING phosphorylation in MCF7 cells (Fig. 8A), despite a reported requirement of STING S366 phosphorylation for STING-mediated IRF3 activation by TBK1 (51-53). We confirmed IRF3 phosphorylation in MCF7 cells in response to dsDNA and also found an increase in overall IRF3 expression (Fig. S3). Moreover, IFN-β was robustly upregulated by ∼ 1000 fold in MCF7 cells (Fig. 8B), while IFN-β upregulation in HeLa cells, in contrast, was an order of magnitude less (Fig. 8C), despite robust STING phosphorylation and a greater increase in CCL22 in HeLa cells. Interestingly, dsDNA did not produce a detectable increase in IFN-β in CCL-2 HeLa cells from ATCC (Fig. 8B), notwithstanding detectable levels of STING phosphorylation (Fig. 7D).

**Figure 8.**
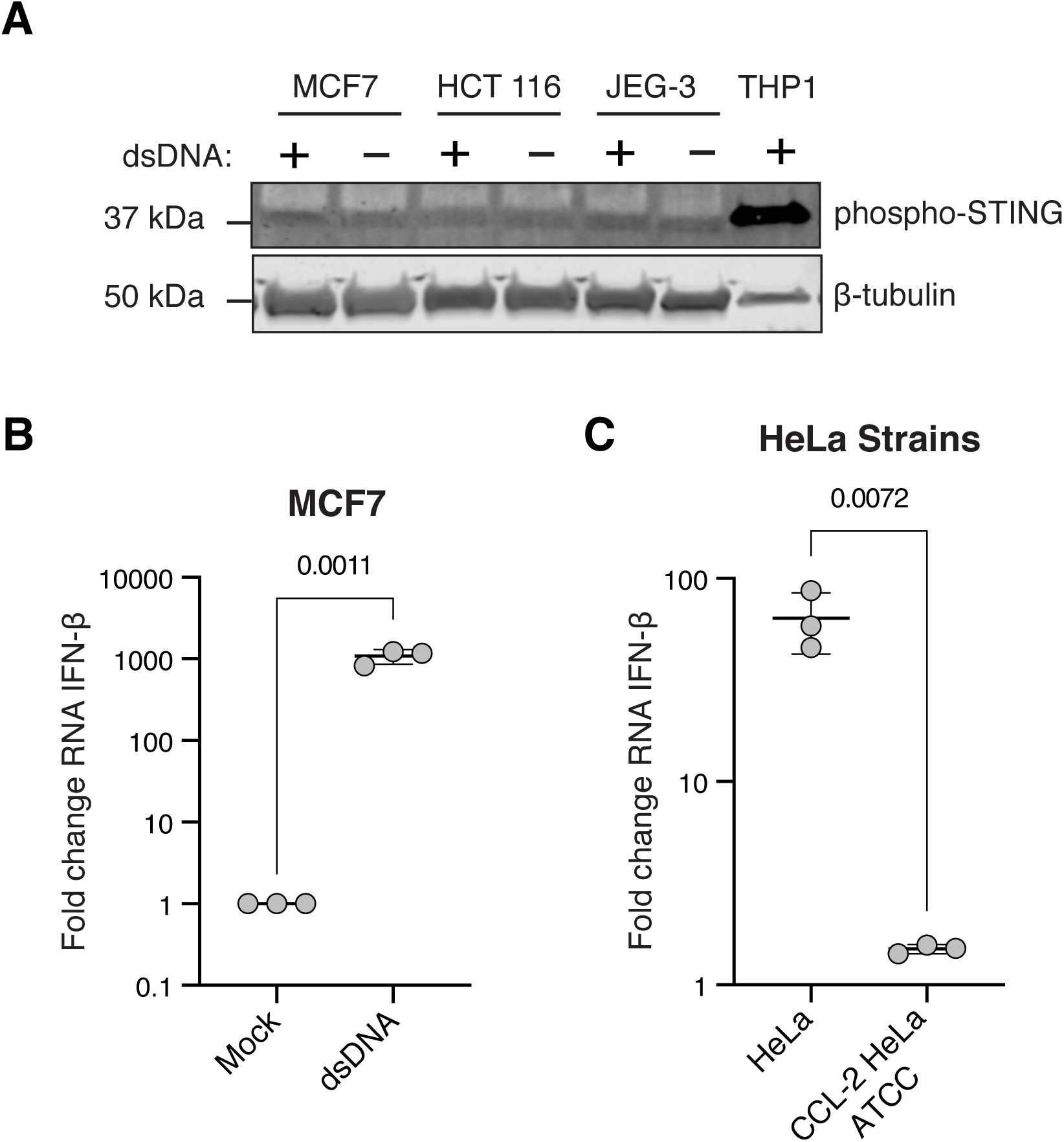
CCL22 is increased despite lack of STING S366 phosphorylation. *A*, Cells were untreated (UN) or transfected with dsDNA (2 μg/mL) using TransfeX at a 1:4 ratio (MCF7) or *Trans*IT-LT1 (HCT 116, 1:4 ratio; JEG-3, 1:3 ratio). Cells were harvested 48 hours after transfection. Lysates (60 μg) and THP1 positive control (15 μg) were separated with SDS-PAGE and probed for phospho-STING (S366) and beta-tubulin. The image shown is representative of at least three independent experiments.

## Discussion

A greater understanding of processes contributing to tumor immune evasion is critical to improving immunotherapy outcomes for solid tumors. A primary mechanism of evasion is recruitment of Tregs to the tumor environment, a process mediated by CCL22 binding to the CCR4 receptor (1-4). We have shown here that dsDNA can robustly upregulate expression of CCL22 in some cancer cells, and that IRF3 is a key regulator of CCL22 expression.

The role of nucleic acid sensing in innate immunity is well-established with respect to pathogen-associated molecular patterns such as double-stranded viral RNA, but its role in anti-tumor immunity in response to self DNA is less completely understood. Leakage of nuclear genomic DNA due to chromosomal instability, DNA released from damaged mitochondria, and re-activation of LINE-1 retrotransposons (46-49,reviewed in 50) can contribute to cGAS activation. Given that each of these are common findings in cancer cells, CCL22 upregulation in response to activated cGAS-STING may be a contributing factor in immune evasion, irrespective of pharmaceutical STING activation.

Multiple questions remain however regarding the regulation and effects of CCL22 in cancer. Previous studies evince a complex CCL22 regulatory landscape that appears to be both species-dependent and cell-type specific. Given the apparent cell-type specificity, one of the more pressing questions warranting further study is whether IRF3 activation is a widespread mechanism of CCL22 upregulation across multiple types of epithelial cancers or only a small subset, as this information may be useful in determining which cancers might have a higher risk of STING-mediated immunosuppression. Our finding that dsDNA did not elicit STING phosphorylation on S366 in MCF7 cells yet nevertheless resulted in IRF3 phosphorylation and upregulation of IFN-β may suggest that other DNA sensing pathways are also involved. Of note, although a majority of studies identified STING S366 phosphorylation as a requirement for dsDNA-mediated activation of IRF3, one study reported that S366 phosphorylation can prevent IRF3 binding to STING (77,reviewed in 78). The relevance of these seemingly opposing functions of STING S366 phosphorylation with respect to dsDNA-mediated upregulation of CCL22 in MCF7 cells remains to be elucidated.

Another issue is whether different STING agonists differentially affect CCL22 expression. In HeLa cells, dsDNA induced higher levels of CCL22 than the STING agonist 2’3’-cGAM(PS)_2_(Rp/Sp), but CCL22 was equivalently induced by each stimulus in MCF7 cells. Different STING agonists can result in distinct functional outcomes (79,reviewed in 80,81), and more work is needed to elucidate how different modes of activation influence CCL22 expression.

It is also interesting to note that STING-mediated immune suppression has been associated with increased indoleamine 2,3-dioxygenase (IDO) expression (54,82), which in turn is reported to be responsive to increased CCL22 (83,84). It was also reported that poly(dA:dT), a synthetic analog of B-DNA that activates DNA sensors as well as RIG-1 via RNA polymerase III (85,reviewed in 86), increased both IDO and CCL22 in some head and neck cancer cells, and that cJUN contributed to CCL22 expression upon direct STING activation (87). These findings highlight the need for a better understanding of factors contributing to CCL22 regulation in cancer cells.

The profound differences observed between the two HeLa cell strains with respect to CCL22 upregulation illustrates that tumor cells can gain, or lose, the capacity to upregulate CCL22 in response to dsDNA. Cancer cell lines, particularly HeLa cells, have been well-documented to evolve in culture in response to myriad selection pressures that vary between laboratories (88). It is unknown whether the different responses of these two strains to dsDNA arose from a single, large-effect mutation or multiple smaller-effect mutations that collectively produced a large phenotypic change, but such evolution and clonal expansion *in vivo* could conceivably contribute to acquired immune evasion. Further comparisons of these two strains may be useful for identifying factors regulating CCL22 in response to dsDNA and STING activation.

Finally, the exact role of IFN-β on CCL22 expression remains to be determined. On one hand, our finding that upregulation of CCL22 occurred only in cells with a coincident increase in IFN-β is consistent with a direct effect of IFN-β, but not indicative. Arguing against a direct effect, previous reports have shown that type I interferons inhibit CCL22 (17,89), and we found very little effect on CCL22 from conditioned media (unpublished data). Determining whether IFN-β and CCL22 are controlled by bifurcated pathways downstream of IRF3 in cancer cells could reveal whether CCL22 might lend itself to independent pharmacological inhibition without affecting IFN-β.

## Experimental procedures

### Cell culture and reagents

Cells were maintained in DMEM high glucose with GlutaMAX and pyruvate (Gibco, cat. 10569010) with 10% FBS and 1x antibiotic-antimycotic (100 units/mL penicillin, 100 ug/mL streptomycin, 0.25 ug/mL Amphotericin B; Gibco, cat. 15240096) in a humidified incubator at 37°C and 5% CO_2_. Two strains of HeLa cervical adenocarcinoma cells were used: one line was a kind gift from Dr. Anthony Furano at the National Institute of Diabetes and Digestive and Kidney Diseases, originally gifted from the late Dr. Haig Kazazian and known in the LINE-1 field as HeLa-JMV; the other strain was purchased from ATCC and designated herein as “CCL-2 HeLa ATCC”. Additional epithelial cancer cell lines purchased from ATCC include MCF7 (mammary gland adenocarcinoma), JEG-3 (placental choriocarcinoma), and HCT 116 (colorectal carcinoma). THP-1 monocytes were also purchased from ATCC. The STING agonist 2’3’-cGAM(PS)_2_(Rp/Sp) (Invivogen, cat. tlrl-nacga2srs) and IRF3 inhibitor MRT67307 (Invivogen, cat. inh-mrt) were each diluted with the companion vial of sterile, endotoxin-free LAL water and used at concentrations indicated in figure legends. STING agonists, immunostimulatory dsDNA (pcDNA3.1(+)puro) and CpG-free dsDNA (pCpGfree-mcs, Invivogen) were transfected using Opti-MEM (Gibco, cat. 31-985-062) as the DNA diluent and either *Trans*IT-LT1 (Mirus, cat. MIR 2300), TransfeX (ATCC, cat. ACS-4005), or *Trans*IT-X2 (Mirus, cat. MIR 6000) based on transfection optimization experiments for each cell line, with the reagent and ratios indicated in figure legends. All cell lines were monitored for mycoplasma using the LookOut mycoplasma PCR detection kit (Sigma-Aldrich, cat. MP0035). THP-1 positive controls for phospho-STING (S366) were generated by plating THP-1 cells in RPMI 1640 media supplemented with 10% FBS and 80 nM PMA for 48 hours to promote differentiation to macrophages before transfecting with pcDNA3.1 (3 μg/mL) using Lipofectamine 3000 at a 1:1 ratio; cells were harvested 4 hours post transfection in 3% SDS, 25 mM Tris-HCl pH 7.4, and 0.5 mM EDTA supplemented with HALT protease and phosphatase inhibitor cocktail at 3X (ThermoFisher, cat. 78440).

### Plasmids

pcDNA3.1(+)-neo was obtained from ThermoFisher (Invitrogen cat. V790-20). pcDNA3.1(+)-neo-IRF3-5D was created using the wild-type template Human V5-IRF3-pcDNA3, a gift from Saumen Sarkar (Addgene plasmid # 32713; http://n2t.net/addgene:32713; RRID:Addgene_32713, (90)) and the Q5 site-directed mutagenesis kit (NEB, cat. E0554S) with the forward and reverse primers 5’-CTCGATCTCGACGATGACCAGTACAAGGCCTAC and 5’-TGGGTGGTCGTTGTCAATGTGCAGGTCCACAGT respectively; mutations were confirmed with sequencing. pCpGf-Bsr-GFP was constructed by PCR-amplifying GFP obtained from pSELECT-zeo-GFPBsr (Invivogen, cat. psetz-zgfpbsr) with a Kozak sequence encoded on the forward primer, then inserting it into pCpGfree-vitroBmcs (Invivogen, cat. pcpgvtb-mcsg2) digested with BglII and ApaL1; insertion was confirmed with sequencing. pcDNA3.1(+)puro was previously described (91). pCpGfree-mcs was obtained from Invivogen (cat. pcpgf-mcs). Lentiviral plasmids included the packaging plasmid psPAX2 (Addgene plasmid # 12260; http://n2t.net/addgene:12260; RRID:Addgene_12260) and the envelope plasmid pMD2.G (Addgene plasmid # 12259; http://n2t.net/addgene:12259; RRID:Addgene_12259), both gifts from Didier Trono.

All RNAi lentiviral expression vectors were pLKO.1-puro, version 1, from The RNAi Consortium (TRC) library collection: control shRNA (Sigma-Aldrich, cat. SHC002); RELA/p65 shRNA-1 (Sigma-Aldrich, cat. TRCN0000014687); RELA/p65 shRNA-2 (Sigma-Aldrich, cat. TRCN0000014684); IRF3 shRNA-1 (Sigma-Aldrich, cat. TRCN0000005921); and IRF3 shRNA-2 (Sigma-Aldrich, cat. TRCN0000005923). DNA for experiments was obtained using endotoxin-free plasmid purification kits (NucleoBond Xtra Midi EF, Takara, cat. 740420.10 or Qiagen EndoFree Plasmid Maxi Kit, cat. 12362); concentration and purity were assessed with spectrophotometry and agarose electrophoresis.

### Lentiviral transductions

HEK293 cells were seeded at 9 × 10^5^ cells per well in 6-well plates and transfected 24 hours later with 1 ug total DNA containing psPAX2, pMD2.G, and each pLKO.1-puro RNAi expression plasmid at a 1:0.25:0.75 ratio, respectively, with *Trans*IT-X2 used at a 1:2 ratio of ug DNA to uL X2. HeLa cells were plated 24 hours prior to transduction at 4 × 10^5^ cells per well in 6-well plates. Media were collected from HEK293 cells 48 hours after transfection and used for transductions in a final concentration of 8 ug/mL polybrene (Sigma-Aldrich, cat. TR-1003). Selection and maintenance of transduced cells was achieved with 5 ug/mL puromycin (Gibco, cat. A1113803) begun 24 hours after application of lentiviral-containing media.

### Live-cell imaging

Live cells were imaged for GFP expression using an EVOS FLoid imaging system. Images were processed using Image J.

### Cell lysis, immunoblots, and antibodies

Cells were lysed for immunoblotting with 3% SDS, 25 mM Tris-HCl pH 7.4, and 0.5 mM EDTA supplemented with 3X HALT protease and phosphatase inhibitor cocktail (ThermoFisher, cat. 78440). Lysates were homogenized with QIAshredder columns (Qiagen, cat. 79656), and protein concentration was determined using the BioRad DC Protein Assay (cat. 5000112). Transfers were performed using either a wet tank or semi-dry system (BioRad Trans-Blot Turbo Transfer System) and low-fluorescent PVDF membranes (BioRad, cat. 1704274 or Millipore Immobilon-FL, cat. IPFL10100). Primary antibodies to the following human proteins were used: phospho-STING (S366, Cell Signaling Technology, cat. 19781S); RELA/p65 (R&D, cat. AF5078-SP); phospho-RELA/p65 (S536, R&D, cat. MAB72261-SP); IRF3 (R&D, cat. AF4019-SP); phospho-IRF3 (S386, Cell Signaling Technology, cat. 37829T); and beta-tubulin loading control (Abcam, cat. ab6046). Fluorescent secondary antibodies were IRDye 680RD and IRDye 800CW (Li-Cor). Blots were imaged with an infrared Li-Cor Odyssey CLx Imager and processed using Image Studio (Li-Cor).

### ELISA

CCL22 protein was measured in media harvested from cells using the MDC human ELISA kit (ABCAM, cat. ab100591) per manufacturer’s instructions.

### RNA purification and RT-qPCR

RNA was isolated by lysing cells in TRIzol reagent (Invitrogen, cat. 15596026) followed by column purification and on-column DNase digestion according to the TRIzol two-step protocol from the Monarch total RNA miniprep kit (NEB, cat. T2010S) or PureLink RNA mini kit (Ambion, cat. 12183018A and 12185010). Quality and concentration were assessed with spectrophotometry. First-strand cDNA synthesis was performed using LunaScript RT SuperMix Kit (NEB, cat. E3010S). qPCR was performed on a QuantStudio 3 or StepOnePlus real-time PCR machine using TaqMan Fast Advanced Master Mix (Applied Biosystems, cat. A44359) and the following TaqMan Assays (assay ID in parenthesis following gene name): CCL22 (Hs00171080); RELA/p65 (Hs00153294_m1); IRF3 (Hs01547283_m1); GAPDH (Hs02786624_g1); beta actin (Hs99999903), 18s (Hs99999901). Data were analyzed using the delta-delta Ct method; housekeeping genes were averaged using the geometric mean.

## Figure Legends

**Supplementary Figure 1.**
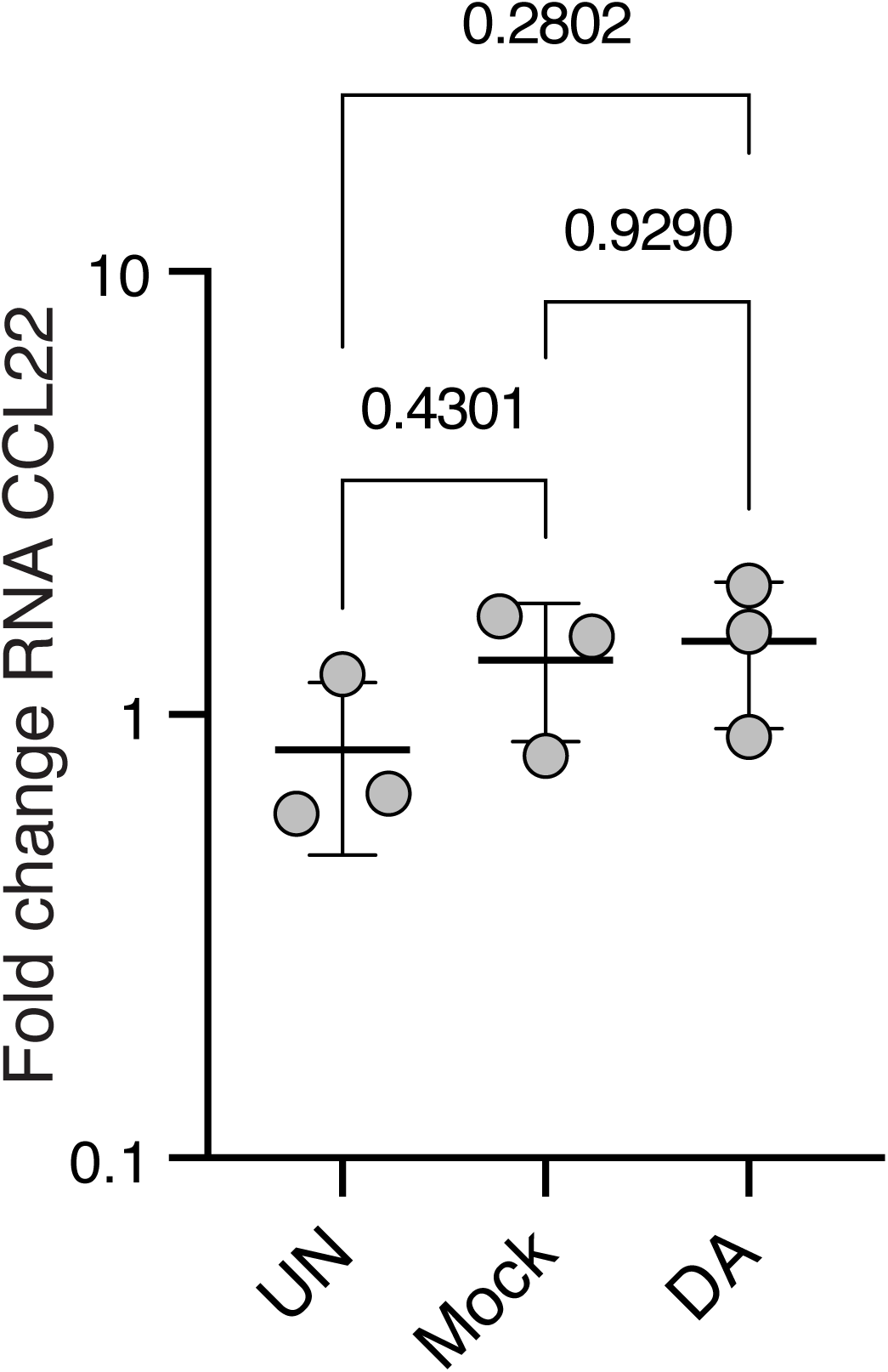
DNA without transfection does not increase CCL22 expression. HeLa cells were untreated (UN) or treated with a mock control containing *Trans*IT-LT1 transfection reagent only or dsDNA alone (DA, 2 μg/mL) without transfection reagent and harvested 48 hours after transfection. Resulting levels of CCL22 mRNA are shown. Each data point represents an independent experiment with values derived from three technical replicates. Significance testing was performed with a one-way ANOVA and Tukey’s pairwise comparison; error bars represent standard deviations.

**Supplementary Figure 2.**
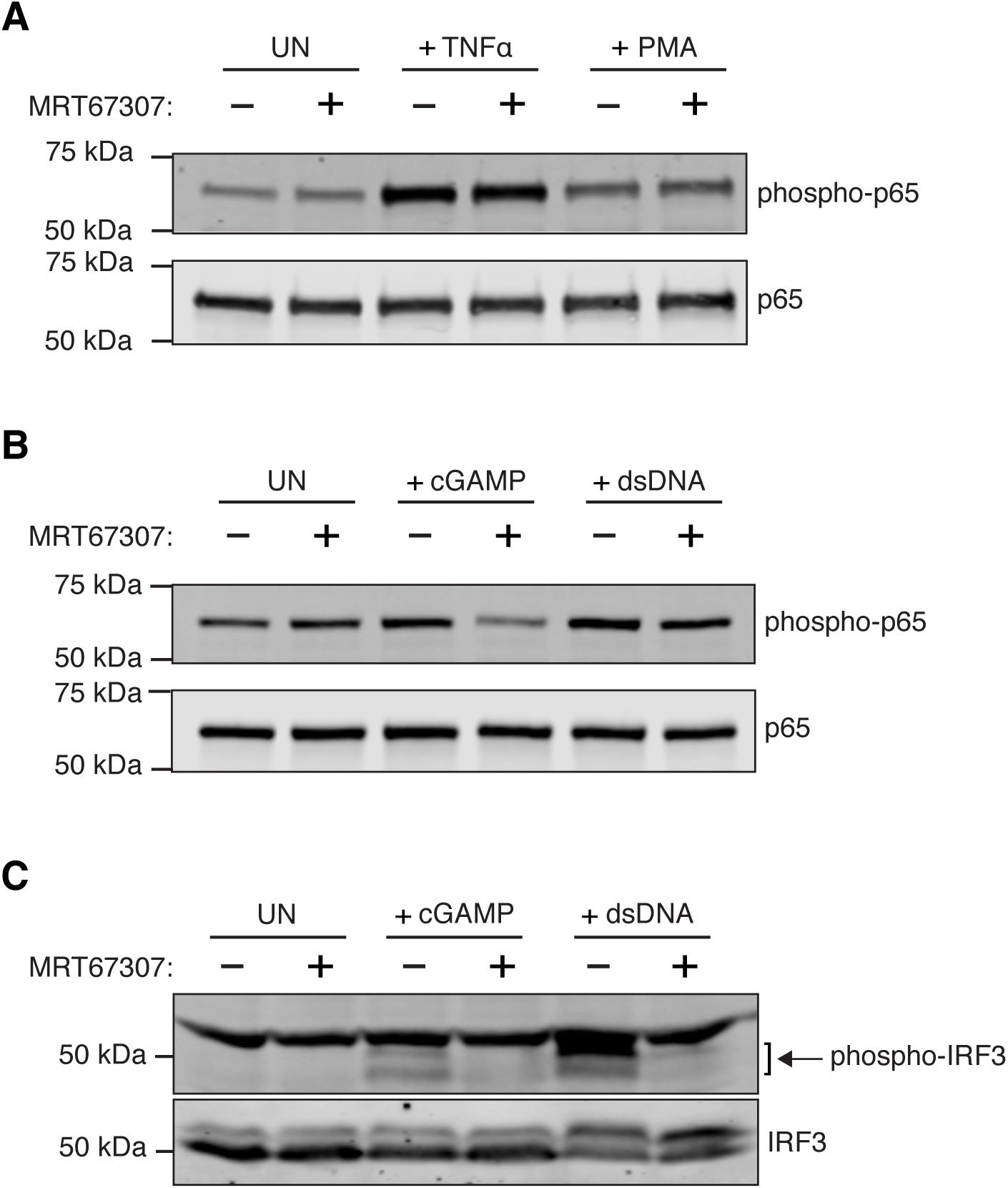
The TBK1/IKKe inhibitor MRT67307 reduces phosphorylation on both IRF3 and RELA/p65. *A*, HeLa cells were treated with a mock water control or 0.6 μM of the TBK1/IKKe inhibitor MRT67307 for 1.5 hours, then treated with TNFα (8 ng/mL) or PMA (10 ng/mL), or untreated (UN) and harvested after 5 minutes (TNFα) or 20 minutes (PMA and UN). Lysates (25 μg) were resolved by SDS-PAGE PAGE and probed for phospho-p65 (S536) and p65. The image shown is representative of at least two independent experiments. *B-C*, HeLa cells were treated with a mock water control or 0.6 μM of the TBK1/IKKe inhibitor MRT67307 for 1.5 hours, then transfected with 10 μM 2’3’-cGAM(PS)_2_(Rp/Sp) and *Trans*IT-LT1 at a 1:1 μg/μL ratio, or dsDNA (2 μg/mL) and *Trans*IT-LT1 at a 1:2 μg/μL ratio, or untreated (UN). Cells were harvested 5 hours after treatment, and lysates were resolved with SDS-PAGE and probed for p-p65 (S536) or p65 as in *A* (25 μg lysate loaded) or for p-IRF3 (S386) and IRF3 (100 μg lysate loaded). Image shown is representative of at least two independent experiments.

**Supplementary Figure 3.**
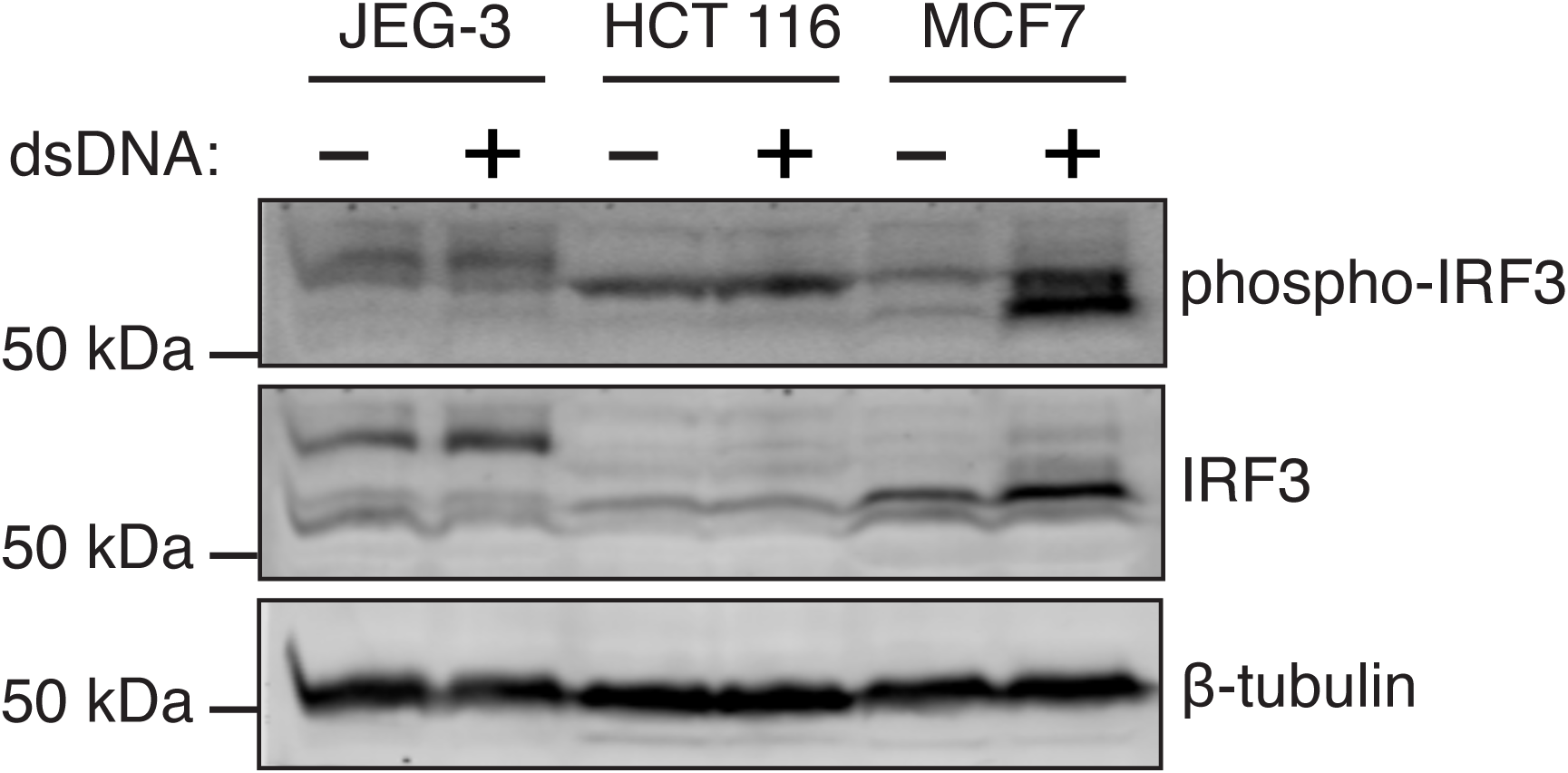
Double-stranded DNA induces IRF3 phosphorylation in MCF7 cells. Cells were untreated (UN) or transfected with dsDNA (2 μg/mL) using *Trans*IT-LT1 (JEG-3, 1:3 ratio; HCT 116, 1:4 ratio) or TransfeX (MCF7, 1:4 ratio). Cells were harvested 48 hours after transfection. Lysates (100 μg) were separated with SDS-PAGE and probed for phospho-IRF3 (S386), IRF3, and beta-tubulin. The image shown is representative of at least two independent experiments.

## Data availability

All data were reported in figures.

## Acknowledgments

We wish to thank Dr. James Drummond and Dr. Robert McKallip for their helpful suggestions during the course of this study. We also thank Dr. Richard McCann for his helpful reading of the manuscript. We would also like to thank Dr. Anthony Furano and the late Dr. Haig Kazazian for the kind gift of HeLa cells.

## Funding and additional information

This work was supported by startup funds and a Dean’s Seed Grant from Mercer University School of Medicine, Macon, Georgia.

## Abbreviations

ATCC: American Type Culture Collection
CDN: cyclic dinucleotide
cGAS: Cyclic GMP-AMP synthase
dsDNA: Double-stranded DNA
GFP: Green fluorescent protein
IFN-γ: Interferon gamma
IKKε: I-kappa-B kinase (IKK) epsilon
IRF3: Interferon regulatory factor 3
MDC: Macrophage derived chemokine
NF-κB: Nuclear factor kappa-light-chain-enhancer of activated B cells
PMA: Phorbol 12-myristate 13-acetate
shRNA: Short hairpin RNA
STING: Stimulator of interferon genes
TBK1: TANK-binding kinase 1
TLR: Toll-like receptor
TME: Tumor microenvironment
TNFα: Tumor necrosis factor alpha
Treg: Regulatory T cell

## References

1. Sugiyama, D., Nishikawa, H., Maeda, Y., Nishioka, M., Tanemura, A., Katayama, I., Ezoe, S., Kanakura, Y., Sato, E., Fukumori, Y., Karbach, J., Jager, E., and Sakaguchi, S. (2013) Anti-CCR4 mAb selectively depletes effector-type FoxP3+CD4+ regulatory T cells, evoking antitumor immune responses in humans. Proc Natl Acad Sci U S A 110, 17945–17950

2. Yoshie, O. (2021) CCR4 as a Therapeutic Target for Cancer Immunotherapy. Cancers (Basel) 13

3. Yoshie, O., and Matsushima, K. (2015) CCR4 and its ligands: from bench to bedside. Int Immunol 27, 11–20

4. Nishikawa, H., and Sakaguchi, S. (2010) Regulatory T cells in tumor immunity. Int J Cancer 127, 759–767

5. Marshall, L. A., Marubayashi, S., Jorapur, A., Jacobson, S., Zibinsky, M., Robles, O., Hu, D. X., Jackson, J. J., Pookot, D., Sanchez, J., Brovarney, M., Wadsworth, A., Chian, D., Wustrow, D., Kassner, P. D., Cutler, G., Wong, B., Brockstedt, D. G., and Talay, O. (2020) Tumors establish resistance to immunotherapy by regulating T(reg) recruitment via CCR4. J Immunother Cancer 8

6. Li, Z. Q., Wang, H. Y., Zeng, Q. L., Yan, J. Y., Hu, Y. S., Li, H., and Yu, Z. J. (2020) p65/miR-23a/CCL22 axis regulated regulatory T cells recruitment in hepatitis B virus positive hepatocellular carcinoma. Cancer Med 9, 711–723

7. Wiedemann, G. M., Röhrle, N., Makeschin, M. C., Fesseler, J., Endres, S., Mayr, D., and Anz, D. (2019) Peritumoural CCL1 and CCL22 expressing cells in hepatocellular carcinomas shape the tumour immune infiltrate. Pathology 51, 586–592

8. Wang, Q., Schmoeckel, E., Kost, B. P., Kuhn, C., Vattai, A., Vilsmaier, T., Mahner, S., Mayr, D., Jeschke, U., and Heidegger, H. H. (2019) Higher CCL22+ Cell Infiltration is Associated with Poor Prognosis in Cervical Cancer Patients. Cancers (Basel) 11

9. Wang, D., Yang, L., Yue, D., Cao, L., Li, L., Wang, D., Ping, Y., Shen, Z., Zheng, Y., Wang, L., and Zhang, Y. (2019) Macrophage-derived CCL22 promotes an immunosuppressive tumor microenvironment via IL-8 in malignant pleural effusion. Cancer Lett 452, 244–253

10. Rapp, M., Wintergerst, M. W. M., Kunz, W. G., Vetter, V. K., Knott, M. M. L., Lisowski, D., Haubner, S., Moder, S., Thaler, R., Eiber, S., Meyer, B., Röhrle, N., Piseddu, I., Grassmann, S., Layritz, P., Kühnemuth, B., Stutte, S., Bourquin, C., von Andrian, U. H., Endres, S., and Anz, D. (2019) CCL22 controls immunity by promoting regulatory T cell communication with dendritic cells in lymph nodes. J Exp Med 216, 1170–1181

11. Jackson, J. J., Ketcham, J. M., Younai, A., Abraham, B., Biannic, B., Beck, H. P., Bui, M. H. T., Chian, D., Cutler, G., Diokno, R., Hu, D. X., Jacobson, S., Karbarz, E., Kassner, P. D., Marshall, L., McKinnell, J., Meleza, C., Okal, A., Pookot, D., Reilly, M. K., Robles, O., Shunatona, H. P., Talay, O., Walker, J. R., Wadsworth, A., Wustrow, D. J., and Zibinsky, M. (2019) Discovery of a Potent and Selective CCR4 Antagonist That Inhibits T(reg) Trafficking into the Tumor Microenvironment. J Med Chem 62, 6190–6213

12. Wei, Y., Wang, T., Song, H., Tian, L., Lyu, G., Zhao, L., and Xue, Y. (2017) C-C motif chemokine 22 ligand (CCL22) concentrations in sera of gastric cancer patients are related to peritoneal metastasis and predict recurrence within one year after radical gastrectomy. J Surg Res 211, 266–278

13. Wiedemann, G. M., Knott, M. M., Vetter, V. K., Rapp, M., Haubner, S., Fesseler, J., Kühnemuth, B., Layritz, P., Thaler, R., Kruger, S., Ormanns, S., Mayr, D., Endres, S., and Anz, D. (2016) Cancer cell-derived IL-1α induces CCL22 and the recruitment of regulatory T cells. Oncoimmunology 5, e1175794

14. Klarquist, J., Tobin, K., Farhangi Oskuei, P., Henning, S. W., Fernandez, M. F., Dellacecca, E. R., Navarro, F. C., Eby, J. M., Chatterjee, S., Mehrotra, S., Clark, J. I., and Le Poole, I. C. (2016) Ccl22 Diverts T Regulatory Cells and Controls the Growth of Melanoma. Cancer Res 76, 6230–6240

15. Chang, D. K., Peterson, E., Sun, J., Goudie, C., Drapkin, R. I., Liu, J. F., Matulonis, U., Zhu, Q., and Marasco, W. A. (2016) Anti-CCR4 monoclonal antibody enhances antitumor immunity by modulating tumor-infiltrating Tregs in an ovarian cancer xenograft humanized mouse model. Oncoimmunology 5, e1090075

16. Jafarzadeh, A., Fooladseresht, H., Minaee, K., Bazrafshani, M. R., Khosravimashizi, A., Nemati, M., Mohammadizadeh, M., Mohammadi, M. M., and Ghaderi, A. (2015) Higher circulating levels of chemokine CCL22 in patients with breast cancer: evaluation of the influences of tumor stage and chemokine gene polymorphism. Tumour Biol 36, 1163–1171

17. Anz, D., Rapp, M., Eiber, S., Koelzer, V. H., Thaler, R., Haubner, S., Knott, M., Nagel, S., Golic, M., Wiedemann, G. M., Bauernfeind, F., Wurzenberger, C., Hornung, V., Scholz, C., Mayr, D., Rothenfusser, S., Endres, S., and Bourquin, C. (2015) Suppression of intratumoral CCL22 by type i interferon inhibits migration of regulatory T cells and blocks cancer progression. Cancer Res 75, 4483–4493

18. Li, Y. Q., Liu, F. F., Zhang, X. M., Guo, X. J., Ren, M. J., and Fu, L. (2013) Tumor secretion of CCL22 activates intratumoral Treg infiltration and is independent prognostic predictor of breast cancer. PLoS One 8, e76379

19. Chang, D. K., Sui, J., Geng, S., Muvaffak, A., Bai, M., Fuhlbrigge, R. C., Lo, A., Yammanuru, A., Hubbard, L., Sheehan, J., Campbell, J. J., Zhu, Q., Kupper, T. S., and Marasco, W. A. (2012) Humanization of an anti-CCR4 antibody that kills cutaneous T-cell lymphoma cells and abrogates suppression by T-regulatory cells. Mol Cancer Ther 11, 2451–2461

20. Faget, J., Biota, C., Bachelot, T., Gobert, M., Treilleux, I., Goutagny, N., Durand, I., Léon-Goddard, S., Blay, J. Y., Caux, C., and Ménétrier-Caux, C. (2011) Early detection of tumor cells by innate immune cells leads to T(reg) recruitment through CCL22 production by tumor cells. Cancer Res 71, 6143–6152

21. Maruyama, T., Kono, K., Izawa, S., Mizukami, Y., Kawaguchi, Y., Mimura, K., Watanabe, M., and Fujii, H. (2010) CCL17 and CCL22 chemokines within tumor microenvironment are related to infiltration of regulatory T cells in esophageal squamous cell carcinoma. Dis Esophagus 23, 422–429

22. Qin, X. J., Shi, H. Z., Deng, J. M., Liang, Q. L., Jiang, J., and Ye, Z. J. (2009) CCL22 recruits CD4-positive CD25-positive regulatory T cells into malignant pleural effusion. Clinical cancer research : an official journal of the American Association for Cancer Research 15, 2231–2237

23. Mailloux, A. W., and Young, M. R. (2009) NK-dependent increases in CCL22 secretion selectively recruits regulatory T cells to the tumor microenvironment. J Immunol 182, 2753–2765

24. Gobert, M., Treilleux, I., Bendriss-Vermare, N., Bachelot, T., Goddard-Leon, S., Arfi, V., Biota, C., Doffin, A. C., Durand, I., Olive, D., Perez, S., Pasqual, N., Faure, C., Ray-Coquard, I., Puisieux, A., Caux, C., Blay, J. Y., and Ménétrier-Caux, C. (2009) Regulatory T cells recruited through CCL22/CCR4 are selectively activated in lymphoid infiltrates surrounding primary breast tumors and lead to an adverse clinical outcome. Cancer Res 69, 2000–2009

25. Wågsäter, D., Dienus, O., Löfgren, S., Hugander, A., and Dimberg, J. (2008) Quantification of the chemokines CCL17 and CCL22 in human colorectal adenocarcinomas. Mol Med Rep 1, 211–217

26. Mizukami, Y., Kono, K., Kawaguchi, Y., Akaike, H., Kamimura, K., Sugai, H., and Fujii, H. (2008) CCL17 and CCL22 chemokines within tumor microenvironment are related to accumulation of Foxp3+ regulatory T cells in gastric cancer. Int J Cancer 122, 2286–2293

27. Tanchot, C., Terme, M., Pere, H., Tran, T., Benhamouda, N., Strioga, M., Banissi, C., Galluzzi, L., Kroemer, G., and Tartour, E. (2013) Tumor-infiltrating regulatory T cells: phenotype, role, mechanism of expansion in situ and clinical significance. Cancer Microenviron 6, 147–157

28. Watanabe, Y., Katou, F., Ohtani, H., Nakayama, T., Yoshie, O., and Hashimoto, K. (2010) Tumor-infiltrating lymphocytes, particularly the balance between CD8(+) T cells and CCR4(+) regulatory T cells, affect the survival of patients with oral squamous cell carcinoma. Oral Surg Oral Med Oral Pathol Oral Radiol Endod 109, 744–752

29. Curiel, T. J., Coukos, G., Zou, L., Alvarez, X., Cheng, P., Mottram, P., Evdemon-Hogan, M., Conejo-Garcia, J. R., Zhang, L., Burow, M., Zhu, Y., Wei, S., Kryczek, I., Daniel, B., Gordon, A., Myers, L., Lackner, A., Disis, M. L., Knutson, K. L., Chen, L., and Zou, W. (2004) Specific recruitment of regulatory T cells in ovarian carcinoma fosters immune privilege and predicts reduced survival. Nat Med 10, 942–949

30. Huang, Y. H., Chang, C. Y., Kuo, Y. Z., Fang, W. Y., Kao, H. Y., Tsai, S. T., and Wu, L. W. (2019) Cancer-associated fibroblast-derived interleukin-1β activates protumor C-C motif chemokine ligand 22 signaling in head and neck cancer. Cancer Sci 110, 2783–2793

31. Kumai, T., Nagato, T., Kobayashi, H., Komabayashi, Y., Ueda, S., Kishibe, K., Ohkuri, T., Takahara, M., Celis, E., and Harabuchi, Y. (2015) CCL17 and CCL22/CCR4 signaling is a strong candidate for novel targeted therapy against nasal natural killer/T-cell lymphoma. Cancer Immunol Immunother 64, 697–705

32. Ménétrier-Caux, C., Faget, J., Biota, C., Gobert, M., Blay, J. Y., and Caux, C. (2012) Innate immune recognition of breast tumor cells mediates CCL22 secretion favoring Treg recruitment within tumor environment. Oncoimmunology 1, 759–761

33. Tsujikawa, T., Yaguchi, T., Ohmura, G., Ohta, S., Kobayashi, A., Kawamura, N., Fujita, T., Nakano, H., Shimada, T., Takahashi, T., Nakao, R., Yanagisawa, A., Hisa, Y., and Kawakami, Y. (2013) Autocrine and paracrine loops between cancer cells and macrophages promote lymph node metastasis via CCR4/CCL22 in head and neck squamous cell carcinoma. Int J Cancer 132, 2755–2766

34. Yang, P., Li, Q. J., Feng, Y., Zhang, Y., Markowitz, G. J., Ning, S., Deng, Y., Zhao, J., Jiang, S., Yuan, Y., Wang, H. Y., Cheng, S. Q., Xie, D., and Wang, X. F. (2012) TGF-β-miR-34a-CCL22 signaling-induced Treg cell recruitment promotes venous metastases of HBV-positive hepatocellular carcinoma. Cancer Cell 22, 291–303

35. Santulli-Marotto, S., Wheeler, J., Lacy, E. R., Boakye, K., Luongo, J., Wu, S. J., and Ryan, M. (2015) CCL22-specific Antibodies Reveal That Engagement of Two Distinct Binding Domains on CCL22 Is Required for CCR4-mediated Function. Monoclon Antib Immunodiagn Immunother 34, 373–380

36. Ashino, S., Wakita, D., Zhang, Y., Chamoto, K., Kitamura, H., and Nishimura, T. (2008) CpG-ODN inhibits airway inflammation at effector phase through down-regulation of antigen-specific Th2-cell migration into lung. Int Immunol 20, 259–266

37. Fujii-Maeda, S., Kajiwara, K., Ikizawa, K., Shinazawa, M., Yu, B., Koga, T., Furue, M., and Yanagihara, Y. (2004) Reciprocal regulation of thymus and activation-regulated chemokine/macrophage-derived chemokine production by interleukin (IL)-4/IL-13 and interferon-gamma in HaCaT keratinocytes is mediated by alternations in E-cadherin distribution. J Invest Dermatol 122, 20–28

38. Horikawa, T., Nakayama, T., Hikita, I., Yamada, H., Fujisawa, R., Bito, T., Harada, S., Fukunaga, A., Chantry, D., Gray, P. W., Morita, A., Suzuki, R., Tezuka, T., Ichihashi, M., and Yoshie, O. (2002) IFN-gamma-inducible expression of thymus and activation-regulated chemokine/CCL17 and macrophage-derived chemokine/CCL22 in epidermal keratinocytes and their roles in atopic dermatitis. Int Immunol 14, 767–773

39. Xiao, T., Kagami, S., Saeki, H., Sugaya, M., Kakinuma, T., Fujita, H., Yano, S., Mitsui, H., Torii, H., Komine, M., Asahina, A., Nakamura, K., and Tamaki, K. (2003) Both IL-4 and IL-13 inhibit the TNF-alpha and IFN-gamma enhanced MDC production in a human keratinocyte cell line, HaCaT cells. J Dermatol Sci 31, 111–117

40. Fukuda, K., Fujitsu, Y., Seki, K., Kumagai, N., and Nishida, T. (2003) Differential expression of thymus-and activation-regulated chemokine (CCL17) and macrophage-derived chemokine (CCL22) by human fibroblasts from cornea, skin, and lung. J Allergy Clin Immunol 111, 520–526

41. Faffe, D. S., Whitehead, T., Moore, P. E., Baraldo, S., Flynt, L., Bourgeois, K., Panettieri, R. A., and Shore, S. A. (2003) IL-13 and IL-4 promote TARC release in human airway smooth muscle cells: role of IL-4 receptor genotype. Am J Physiol Lung Cell Mol Physiol 285, L907–914

42. Bonecchi, R., Sozzani, S., Stine, J. T., Luini, W., D’Amico, G., Allavena, P., Chantry, D., and Mantovani, A. (1998) Divergent effects of interleukin-4 and interferon-gamma on macrophage-derived chemokine production: an amplification circuit of polarized T helper 2 responses. Blood 92, 2668–2671

43. Galli, G., Chantry, D., Annunziato, F., Romagnani, P., Cosmi, L., Lazzeri, E., Manetti, R., Maggi, E., Gray, P. W., and Romagnani, S. (2000) Macrophage-derived chemokine production by activated human T cells in vitro and in vivo: preferential association with the production of type 2 cytokines. Eur J Immunol 30, 204–210

44. Hopfner, K. P., and Hornung, V. (2020) Molecular mechanisms and cellular functions of cGAS-STING signalling. Nat Rev Mol Cell Biol 21, 501–521

45. Yu, L., and Liu, P. (2021) Cytosolic DNA sensing by cGAS: regulation, function, and human diseases. Signal Transduct Target Ther 6, 170

46. Simon, M., Van Meter, M., Ablaeva, J., Ke, Z., Gonzalez, R. S., Taguchi, T., De Cecco, M., Leonova, K. I., Kogan, V., Helfand, S. L., Neretti, N., Roichman, A., Cohen, H. Y., Meer, M. V., Gladyshev, V. N., Antoch, M. P., Gudkov, A. V., Sedivy, J. M., Seluanov, A., and Gorbunova, V. (2019) LINE1 Derepression in Aged Wild-Type and SIRT6-Deficient Mice Drives Inflammation. Cell Metab 29, 871–885 e875

47. Stetson, D. B., Ko, J. S., Heidmann, T., and Medzhitov, R. (2008) Trex1 prevents cell-intrinsic initiation of autoimmunity. Cell 134, 587–598

48. Thomas, C. A., Tejwani, L., Trujillo, C. A., Negraes, P. D., Herai, R. H., Mesci, P., Macia, A., Crow, Y. J., and Muotri, A. R. (2017) Modeling of TREX1-Dependent Autoimmune Disease using Human Stem Cells Highlights L1 Accumulation as a Source of Neuroinflammation. Cell Stem Cell 21, 319–331 e318

49. De Cecco, M., Ito, T., Petrashen, A. P., Elias, A. E., Skvir, N. J., Criscione, S. W., Caligiana, A., Brocculi, G., Adney, E. M., Boeke, J. D., Le, O., Beausejour, C., Ambati, J., Ambati, K., Simon, M., Seluanov, A., Gorbunova, V., Slagboom, P. E., Helfand, S. L., Neretti, N., and Sedivy, J. M. (2019) Author Correction: L1 drives IFN in senescent cells and promotes age-associated inflammation. Nature 572, E5

50. Decout, A., Katz, J. D., Venkatraman, S., and Ablasser, A. (2021) The cGAS-STING pathway as a therapeutic target in inflammatory diseases. Nat Rev Immunol 21, 548–569

51. Tanaka, Y., and Chen, Z. J. (2012) STING specifies IRF3 phosphorylation by TBK1 in the cytosolic DNA signaling pathway. Sci Signal 5, ra20

52. Liu, S., Cai, X., Wu, J., Cong, Q., Chen, X., Li, T., Du, F., Ren, J., Wu, Y. T., Grishin, N. V., and Chen, Z. J. (2015) Phosphorylation of innate immune adaptor proteins MAVS, STING, and TRIF induces IRF3 activation. Science 347, aaa2630

53. Zhang, C., Shang, G., Gui, X., Zhang, X., Bai, X. C., and Chen, Z. J. (2019) Structural basis of STING binding with and phosphorylation by TBK1. Nature 567, 394–398

54. Lemos, H., Mohamed, E., Huang, L., Ou, R., Pacholczyk, G., Arbab, A. S., Munn, D., and Mellor, A. L. (2016) STING Promotes the Growth of Tumors Characterized by Low Antigenicity via IDO Activation. Cancer Res 76, 2076–2081

55. Kwon, J., and Bakhoum, S. F. (2020) The Cytosolic DNA-Sensing cGAS-STING Pathway in Cancer. Cancer Discov 10, 26–39

56. Jiang, M., Chen, P., Wang, L., Li, W., Chen, B., Liu, Y., Wang, H., Zhao, S., Ye, L., He, Y., and Zhou, C. (2020) cGAS-STING, an important pathway in cancer immunotherapy. J Hematol Oncol 13, 81

57. Bakhoum, S. F., Ngo, B., Laughney, A. M., Cavallo, J. A., Murphy, C. J., Ly, P., Shah, P., Sriram, R. K., Watkins, T. B. K., Taunk, N. K., Duran, M., Pauli, C., Shaw, C., Chadalavada, K., Rajasekhar, V. K., Genovese, G., Venkatesan, S., Birkbak, N. J., McGranahan, N., Lundquist, M., LaPlant, Q., Healey, J. H., Elemento, O., Chung, C. H., Lee, N. Y., Imielenski, M., Nanjangud, G., Pe’er, D., Cleveland, D. W., Powell, S. N., Lammerding, J., Swanton, C., and Cantley, L. C. (2018) Chromosomal instability drives metastasis through a cytosolic DNA response. Nature 553, 467–472

58. Ahn, J., Xia, T., Konno, H., Konno, K., Ruiz, P., and Barber, G. N. (2014) Inflammation-driven carcinogenesis is mediated through STING. Nat Commun 5, 5166

59. Chan, M. P., Onji, M., Fukui, R., Kawane, K., Shibata, T., Saitoh, S., Ohto, U., Shimizu, T., Barber, G. N., and Miyake, K. (2015) DNase II-dependent DNA digestion is required for DNA sensing by TLR9. Nat Commun 6, 5853

60. Miyake, K., Shibata, T., Ohto, U., Shimizu, T., Saitoh, S. I., Fukui, R., and Murakami, Y. (2018) Mechanisms controlling nucleic acid-sensing Toll-like receptors. Int Immunol 30, 43–51

61. Abe, T., and Barber, G. N. (2014) Cytosolic-DNA-mediated, STING-dependent proinflammatory gene induction necessitates canonical NF-kappaB activation through TBK1. J Virol 88, 5328–5341

62. Fang, R., Wang, C., Jiang, Q., Lv, M., Gao, P., Yu, X., Mu, P., Zhang, R., Bi, S., Feng, J. M., and Jiang, Z. (2017) NEMO-IKKbeta Are Essential for IRF3 and NF-kappaB Activation in the cGAS-STING Pathway. J Immunol 199, 3222–3233

63. Balka, K. R., Louis, C., Saunders, T. L., Smith, A. M., Calleja, D. J., D’Silva, D. B., Moghaddas, F., Tailler, M., Lawlor, K. E., Zhan, Y., Burns, C. J., Wicks, I. P., Miner, J. J., Kile, B. T., Masters, S. L., and De Nardo, D. (2020) TBK1 and IKKepsilon Act Redundantly to Mediate STING-Induced NF-kappaB Responses in Myeloid Cells. Cell Rep 31, 107492

64. Yum, S., Li, M., Fang, Y., and Chen, Z. J. (2021) TBK1 recruitment to STING activates both IRF3 and NF-kappaB that mediate immune defense against tumors and viral infections. Proc Natl Acad Sci U S A 118

65. Mattioli, I., Geng, H., Sebald, A., Hodel, M., Bucher, C., Kracht, M., and Schmitz, M. L. (2006) Inducible phosphorylation of NF-kappa B p65 at serine 468 by T cell costimulation is mediated by IKK epsilon. J Biol Chem 281, 6175–6183

66. Christian, F., Smith, E. L., and Carmody, R. J. (2016) The Regulation of NF-kappaB Subunits by Phosphorylation. Cells 5

67. Poole, E., Atkins, E., Nakayama, T., Yoshie, O., Groves, I., Alcami, A., and Sinclair, J. (2008) NF-kappaB-mediated activation of the chemokine CCL22 by the product of the human cytomegalovirus gene UL144 escapes regulation by viral IE86. J Virol 82, 4250–4256

68. Nakayama, T., Hieshima, K., Nagakubo, D., Sato, E., Nakayama, M., Kawa, K., and Yoshie, O. (2004) Selective induction of Th2-attracting chemokines CCL17 and CCL22 in human B cells by latent membrane protein 1 of Epstein-Barr virus. J Virol 78, 1665–1674

69. Sun, J., Sun, J., Song, B., Zhang, L., Shao, Q., Liu, Y., Yuan, D., Zhang, Y., and Qu, X. (2016) Fucoidan inhibits CCL22 production through NF-kappaB pathway in M2 macrophages: a potential therapeutic strategy for cancer. Sci Rep 6, 35855

70. Ghadially, H., Ross, X. L., Kerst, C., Dong, J., Reske-Kunz, A. B., and Ross, R. (2005) Differential regulation of CCL22 gene expression in murine dendritic cells and B cells. J Immunol 174, 5620–5629

71. Qi, X. F., Kim, D. H., Yoon, Y. S., Li, J. H., Song, S. B., Jin, D., Huang, X. Z., Teng, Y. C., and Lee, K. J. (2009) The adenylyl cyclase-cAMP system suppresses TARC/CCL17 and MDC/CCL22 production through p38 MAPK and NF-kappaB in HaCaT keratinocytes. Mol Immunol 46, 1925–1934

72. Kwon, D. J., Bae, Y. S., Ju, S. M., Goh, A. R., Youn, G. S., Choi, S. Y., and Park, J. (2012) Casuarinin suppresses TARC/CCL17 and MDC/CCL22 production via blockade of NF-kappaB and STAT1 activation in HaCaT cells. Biochem Biophys Res Commun 417, 1254–1259

73. Lin, R., Heylbroeck, C., Pitha, P. M., and Hiscott, J. (1998) Virus-dependent phosphorylation of the IRF-3 transcription factor regulates nuclear translocation, transactivation potential, and proteasome-mediated degradation. Mol Cell Biol 18, 2986–2996

74. Lin, R., Mamane, Y., and Hiscott, J. (1999) Structural and functional analysis of interferon regulatory factor 3: localization of the transactivation and autoinhibitory domains. Mol Cell Biol 19, 2465–2474

75. Panne, D., McWhirter, S. M., Maniatis, T., and Harrison, S. C. (2007) Interferon regulatory factor 3 is regulated by a dual phosphorylation-dependent switch. J Biol Chem 282, 22816–22822

76. Schwanke, H., Stempel, M., and Brinkmann, M. M. (2020) Of Keeping and Tipping the Balance: Host Regulation and Viral Modulation of IRF3-Dependent IFNB1 Expression. Viruses 12

77. Li, Y., Wilson, H. L., and Kiss-Toth, E. (2017) Regulating STING in health and disease. J Inflamm (Lond) 14, 11

78. Konno, H., Konno, K., and Barber, G. N. (2013) Cyclic dinucleotides trigger ULK1 (ATG1) phosphorylation of STING to prevent sustained innate immune signaling. Cell 155, 688–698

79. Li, S., Luo, M., Wang, Z., Feng, Q., Wilhelm, J., Wang, X., Li, W., Wang, J., Cholka, A., Fu, Y. X., Sumer, B. D., Yu, H., and Gao, J. (2021) Author Correction: Prolonged activation of innate immune pathways by a polyvalent STING agonist. Nat Biomed Eng 5, 483

80. Cui, X., Zhang, R., Cen, S., and Zhou, J. (2019) STING modulators: Predictive significance in drug discovery. Eur J Med Chem 182, 111591

81. Motedayen Aval, L., Pease, J. E., Sharma, R., and Pinato, D. J. (2020) Challenges and Opportunities in the Clinical Development of STING Agonists for Cancer Immunotherapy. J Clin Med 9

82. Lemos, H., Ou, R., McCardle, C., Lin, Y., Calver, J., Minett, J., Chadli, A., Huang, L., and Mellor, A. L. (2020) Overcoming resistance to STING agonist therapy to incite durable protective antitumor immunity. J Immunother Cancer 8

83. Onodera, T., Jang, M. H., Guo, Z., Yamasaki, M., Hirata, T., Bai, Z., Tsuji, N. M., Nagakubo, D., Yoshie, O., Sakaguchi, S., Takikawa, O., and Miyasaka, M. (2009) Constitutive expression of IDO by dendritic cells of mesenteric lymph nodes: functional involvement of the CTLA-4/B7 and CCL22/CCR4 interactions. J Immunol 183, 5608–5614

84. Bischoff, L., Alvarez, S., Dai, D. L., Soukhatcheva, G., Orban, P. C., and Verchere, C. B. (2015) Cellular mechanisms of CCL22-mediated attenuation of autoimmune diabetes. J Immunol 194, 3054–3064

85. Chiu, Y. H., Macmillan, J. B., and Chen, Z. J. (2009) RNA polymerase III detects cytosolic DNA and induces type I interferons through the RIG-I pathway. Cell 138, 576–591

86. Unterholzner, L. (2013) The interferon response to intracellular DNA: why so many receptors? Immunobiology 218, 1312–1321

87. Liang, D., Xiao-Feng, H., Guan-Jun, D., Er-Ling, H., Sheng, C., Ting-Ting, W., Qin-Gang, H., Yan-Hong, N., and Ya-Yi, H. (2015) Activated STING enhances Tregs infiltration in the HPV-related carcinogenesis of tongue squamous cells via the c-jun/CCL22 signal. Biochim Biophys Acta 1852, 2494–2503

88. Liu, Y., Mi, Y., Mueller, T., Kreibich, S., Williams, E. G., Van Drogen, A., Borel, C., Frank, M., Germain, P. L., Bludau, I., Mehnert, M., Seifert, M., Emmenlauer, M., Sorg, I., Bezrukov, F., Bena, F. S., Zhou, H., Dehio, C., Testa, G., Saez-Rodriguez, J., Antonarakis, S. E., Hardt, W. D., and Aebersold, R. (2019) Multi-omic measurements of heterogeneity in HeLa cells across laboratories. Nat Biotechnol 37, 314–322

89. Iellem, A., Colantonio, L., Bhakta, S., Sozzani, S., Mantovani, A., Sinigaglia, F., and D’Ambrosio, D. (2000) Inhibition by IL-12 and IFN-alpha of I-309 and macrophage-derived chemokine production upon TCR triggering of human Th1 cells. Eur J Immunol 30, 1030–1039

90. Zhu, J., Smith, K., Hsieh, P. N., Mburu, Y. K., Chattopadhyay, S., Sen, G. C., and Sarkar, S. N. (2010) High-throughput screening for TLR3-IFN regulatory factor 3 signaling pathway modulators identifies several antipsychotic drugs as TLR inhibitors. J Immunol 184, 5768–5776

91. Cook, P. R., Jones, C. E., and Furano, A. V. (2015) Phosphorylation of ORF1p is required for L1 retrotransposition. Proc Natl Acad Sci U S A 112, 4298–4303

